# Comprehensive genomic analysis reveals virulence factors and antibiotic resistance genes in *Pantoea agglomerans* KM1, a potential opportunistic pathogen

**DOI:** 10.1101/2020.09.15.297663

**Authors:** Robin B. Guevarra, Stefan Magez, Eveline Peeters, Mi Sook Chung, Kyung Hyun Kim, Magdalena Radwanska

## Abstract

*Pantoea agglomerans* is a Gram-negative aerobic bacillus causing a wide range of opportunistic infections in humans including septicemia, pneumonia, septic arthritis, wound infections and meningitis. To date, the determinants of virulence, antibiotic resistance, metabolic features conferring survival and host-associated pathogenic potential of this bacterium remain largely underexplored. In this study, we sequenced and assembled the whole-genome of *P. agglomerans* KM1 isolated from kimchi in South Korea. The genome contained one circular chromosome of 4,039,945 bp, 3 mega plasmids, and 2 prophages. The phage-derived genes encoded integrase, lysozyme and terminase. Six CRISPR loci were identified within the bacterial chromosome. Further in-depth analysis showed that the genome contained 13 antibiotic resistance genes conferring resistance to clinically important antibiotics such as penicillin G, bacitracin, rifampicin, vancomycin, and fosfomycin. Genes involved in adaptations to environmental stress were also identified which included factors providing resistance to osmotic lysis, oxidative stress, as well as heat and cold shock. The genomic analysis of virulence factors led to identification of a type VI secretion system, hemolysin, filamentous hemagglutinin, and genes involved in iron uptake and sequestration. Finally, the data provided here show that, the KM1 isolate exerted strong immunostimulatory properties on RAW 264.7 macrophages *in vitr*o. Stimulated cells produced Nitric Oxide (NO) and pro-inflammatory cytokines TNF-α, IL-6 and the anti-inflammatory cytokine IL-10. The upstream signaling for production of TNF-α, IL-6, IL-10, and NO depended on TLR4 and TLR1/2. While production of TNF-α, IL-6 and NO involved solely activation of the NF-κB, IL-10 secretion was largely dependent on NF-κB and to a lesser extent on MAPK Kinases. Taken together, the analysis of the whole-genome and immunostimulatory properties provided in-depth characterization of the *P. agglomerans* KM1 isolate shedding a new light on determinants of virulence that drive its interactions with the environment, other microorganisms and eukaryotic hosts.

## Introduction

Kimchi is a well-known Korean vegetable side dish prepared based on fermentation process conducted by lactic acid bacteria (LAB) [1, 2]. Often homemade, kimchi is traditionally considered as a functional food with potential health benefits including anti-cancer, anti-obesity, anti-oxidant and anti-aging properties [1]. The fermentation process of a Korean Baechu cabbage leads to steadily decreasing pH, reaching 4.5 after approximately one month of fermentation. This process is followed by a subsequent growth of the LAB and elimination of potentially pathogenic bacterial species that are sensitive to low pH [3]. Recently, the interest in consumption of kimchi has increased leading to a global distribution of this traditional Korean side dish. However, the time needed for a complete fermentation to take place is often reduced in favor of a ‘better’ taste, resulting in frequent consumption of a ‘short-term’ fermented kimchi. This, in turn, poses potential health risks as previous publications reported several outbreaks associated with a foodborne pathogen-contaminated kimchi [4, 5]. It was shown that pathogenic *Escherichia coli* and *Salmonella* can survive in kimchi during fermentation, suggesting that the risk for contamination of kimchi with pathogens, in both commercial and homemade preparations, should not be underestimated [6].

*Pantoea agglomerans* (formerly *Enterobacter agglomerans* and *Erwinia herbicola*) is a Gram-negative member of the *Enterobacteriaceae* family that is ubiquitous in nature and is found in a wide variety of environments. *P. agglomerans* grows as epiphyte on plants [7], exists in soil [8], insects [9] various food sources [10] and has been found in clinical samples including pus, sputum, urine, bloodstream, tracheal and joint aspirate [11-14]. Different strains of *P. agglomerans* exert beneficial antibacterial activity against phytopathogens [15], which make them an attractive biocontrol agent [16] and a plant growth promoter [17]. While *P. agglomerans* strains E325 and P10c biocontrol agents are allowed for agricultural use in several countries including USA and New Zealand, it remains to be approved for commercial purposes by the European countries [18]. The lipopolysaccharide (LPS) extracted from *P. agglomerans* is a potent adjuvant in mucosal vaccinations, triggering production of IgGs and IgAs, and inducting TNF-α and IL-6 secretion through activation of a Toll-like receptor 4 (TLR4) [19]. *P. agglomerans* derived LPS has anti-cancer properties in B16 melanoma model in mice [20], and an oral administration of this molecule contributed to prevention of atherosclerosis and hypertension [21].

While *P. agglomerans* may exert some beneficial roles, it can cause life-threatening infections in immunosuppressed individuals, elderly, newborns and infants [12, 22, 23]. For example, clinical isolates containing *P. agglomerans* can cause a wide range of infections such as neonatal sepsis [24], joint infection [11], pneumonia and meningitis [25]. *P. agglomerans* furthermore poses health risks for adults, being responsible for occupational respiratory diseases of workers exposed to dust in factories processing cotton, herbs, grain, wood, and tobacco [22, 26, 27]. Affected individuals suffered from inflammation of the respiratory system and allergic pulmonary disorders [23]. In animal studies, inhaled nanoparticles carrying *P. agglomerans* LPS induced airway inflammation with alveolar macrophages secreting pro-inflammatory cytokines and superoxide anion [28]. Other studies conducted using experimental mouse models showed that ingested *P. agglomerans* was able to colonize the gut, cross the gut barrier to the bloodstream, and cause systemic dissemination to various organs [29]. Taken together, a number of reports demonstrated the ambiguous nature of *P. agglomerans* in relation to possible beneficial or detrimental effects on health. In this context, there is scarce information about possible determinants of virulence leading to pathology of *P. agglomerans* associated diseases.

Advances in genome sequencing allowed for identification of virulence factors, and plant growth promoting determinants in genomes of various species belonging to genus *Pantoea* [7, 17, 30]. In plants, some of the isolated pathogenic stains of *P. agglomerans* are known to cause gall-formation, which contained type III secretion system (T3SS) and its effectors through acquisition of pathogenicity plasmid (pPATH). *P. agglomerans* biopesticide properties are related to the presence of genes encoding antibiotics such as pantocins, herbicolins, microcins, and phenazines, which target for example amino acid biosynthesis genes in *Erwinia amylovora*, the causative agent of fire blight [14].

While most *P. agglomerans* genome studies focused on plant isolates, the determinants of virulence present in clinical isolates remain largely undiscovered. Our current study aimed at filling this knowledge gap based on the analysis of the sequenced genome and tested immuno-properties of the isolated foodborne *P. agglomerans* KM1 strain. We identified genes involved in virulence, antibiotic resistance, adaptations to stress and interactions with other microorganisms and eukaryotic hosts. The study also demonstrated immuno-modulatory properties of *P. agglomerans* KM1 and mechanisms involved in secretion of pro-inflammatory and anti-inflammatory cytokines.

## Materials and Methods

### Isolation and growth conditions

*P. agglomerans* KM1 was isolated from short-term fermented homemade kimchi pH 4.5 delivered to the laboratory. All isolation procedures were conducted in sterile conditions. Blended kimchi leaves and juice samples were filtered through sterile gauze and spun down for 15 seconds to recover liquid fractions. These were plated out onto Luria Bertani (LB) agar plate and incubated overnight at 37°C. From the kimchi juice and leaves, eight and nineteen glistening yellow-pigmented colonies were counted on the 10^−4^ diluted sample, respectively.

### Genomic DNA extraction and 16S rRNA based identification

A single colony of *P. agglomerans* KM1 was grown in LB broth (Conda, Spain) overnight at 37°C. The genomic DNA was isolated using the LaboPass™ Tissue Genomic DNA mini kit (Cosmo Genetech, Seoul, South Korea) according to the manufacturer’s protocol. The identity of the isolate was verified through 16S rRNA gene sequencing. The 16S rRNA gene was amplified from the extracted genomic DNA using 16S universal primers 27F (5’-AGA GTTTGATCMTGGCTCAG-3’) and 1492R (5’-GGTTACCTTGTTACGACTTC-3’) and sequenced using an automated ABI3730XL capillary DNA sequencer (Applied Biosystems, USA) for taxonomic identification at Cosmo Genetech (Seoul, South Korea). The 16S rRNA sequences were confirmed through BLASTn search against the NCBI microbial 16S database.

### Genome sequencing and assembly

The genome of *P. agglomerans* KM1 was sequenced using the Illumina HiSeq 4000 and 2 × 150 bp with an insert size of 350 bp at Macrogen, Inc. (Seoul, South Korea). Libraries were generated from1 µg of genomic DNA using the TruSeq® DNA PCR-free Library Prep Kit (Illumina) according to the manufacturer’s protocol. The quality control of the raw reads was performed using FASTQC v.0.11.9, and quality trimming was done based on FASTQC report using Trimmomatic v.0.39. Obtained quality trimmed reads were used for *de novo* assembly using SPAdes v3.14.1 [31] with default parameters. The final genome assembly was polished with the Illumina reads using Pilon v.1.23 [32]. The quality of the genome assembly was evaluated using QUAST v5.0.2 [33], and the assessment of the genome completeness was performed using BUSCO v4.0.5 [34]. The resulting contigs were reordered using the Mauve Contig Mover algorithm in Mauve v2.4.0 against the closest complete genome of *Pantoea agglomerans* C410P1 as a reference (GenBank accession number CP016889). The whole-genome shotgun project of *P. agglomerans* KM1 was deposited at NCBI GenBank under accession number NZ_JAAVXI000000000.

### Genome annotation

Functional genome annotation of *P. agglomerans* KM1 was conducted using NCBI Prokaryotic Genome Annotation Pipeline (PGAP) [35]. The tRNA genes were identified by tRNAscan-SE v.2.0 [36] and rRNA genes with RNAmmer v.1.2 [37]. The Cluster of Orthologous Groups (COG) functional annotation was performed using RPS-BLAST program against the COG database within the WebMGA server [38]. For Subsystem functional categorization, SEED annotation was used with the SEED viewer within the Rapid Annotations and applying Subsystems Technology (RAST) server v2.0 [39]. The circular genome maps were generated using the GView v1.7 [40].

### Phylogenetic analysis

Phylogenetic trees were constructed based on the average nucleotide identity (ANI) and 16S rRNA gene. The overall similarity between the whole-genome sequences was calculated using the Orthologous Average Nucleotide Identity Tool (OAT) v0.93.1 [41]. Phylogenetic relationships based on the 16S rRNA gene sequences between *P. agglomerans* KM1 and closely related species were determined using the Molecular Evolutionary Genetic Analysis X (MEGA X) software using the neighbor-joining method with 1,000 randomly selected bootstrap replicates [42]. The NCBI GenBank assembly accession numbers of the *Pantoea* genomes used for phylogenetic analysis are the following: *P. agglomerans* KM1 (GCA_012241415.1), *P. agglomerans* C410P1 (GCA_001709315.1), *P. agglomerans* UAEU18 (GCA_010523255.1), *P. agglomerans* Tx10 (GCA_000475055.1), *P. agglomerans* IG1 (GCA_000241285.2), *P. agglomerans* JM1 (GCA_002222515.1), *P. agglomerans* P5 (GCA_002157425.2), *P. agglomerans* TH81 (GCA_010523255.1), *P. agglomerans* L15 (GCA_003860325.1), *P. vagans* MP7 (GCA_000757435.1), *P. vagans* C9-1 (GCA_000148935.1), *P. ananatis* LMG 2665 (GCA_000661975.1) and *Escherichia coli* strain K12 substr. MG1655 (GCA_000005845.2).

### Comparative genomics

The genome synteny analysis was performed using the progressive Mauve algorithm in Mauve v2.3.0 [43] to visualize the genome alignments of the *P. agglomerans* KM1 draft genome with *P. agglomerans* C410P1 as a reference (GenBank accession number CP016889). The Mauve software was used to predict chromosomal rearrangement such as insertion, inversion and translocation. The pan-genome analysis was conducted using the retrieved published *P. agglomerans* complete genomes in the NCBI database including those of C410P1 (GCA_001709315.1), UAEU18 (GCA_010523255.1), TH81 (GCA_003704305.1), and L15 (GCA_003860325.1). Prior to pan-genome analysis, Rapid Prokaryotic Genome Annotation (PROKKA) [44] pipeline was used to annotate the genome sequences with Roary software [45] to create the pan-genome and using the Gview Server for visualization [40].

### Prediction of genomic islands, prophages and CRISPRs

Identification and visualization of putative genomic islands were conducted using IslandPick, IslandPath-DIMOB, and SIGI-HMM prediction methods in IslandViewer4 [46]. Prophages were identified using PHAge Search Tool Enhanced Release (PHASTER) [47]. CRISPRFinder software was used to identify clustered regularly interspaced short palindromic repeats (CRISPRs) [48].

### Biochemical characterization

The isolate was tested for presence of a cytochrome oxidase using the 70439 Oxidase assay (Sigma-Aldrich, MO, USA) according to instruction provided by the manufacturer. The presence of a catalase activity was assessed by exposure of a bacterial colony to 3% hydrogen peroxide (Daeyung, Gyeonggi-do, South Korea). Further biochemical characterization was performed using the Analytical Profile Index (API) 20E test (BioMérieux, France) according to manufacturer’s protocol. The index profile was determined with the online API 20E v5.0 identification software (apiweb.biomerieux.com).

### Antibiotic resistance gene identification and susceptibility testing

Antibiotic resistance genes (ARGs) were predicted using the BLASTn method against the Comprehensive Antibiotic Resistance Database (CARD) [49] using an identity cut-off of 70% and an E-value < 1.0 E^-6^. Antibiotic susceptibility test was done using the Kirby-Bauer disk diffusion method on Mueller Hinton agar according to the Clinical and Laboratory Standards Institute (CLSI). Commercially prepared antibiotic disks used in this study were kanamycin (30 µg), streptomycin (10 µg), imipenem (10 µg), vancomycin (10 µg), ofloxacin (5 µg), ampicillin (30 µg), penicillin G (10 iu), rifampicin (5 µg), bacitracin (10 µg), fosfomycin (50 µg), and chloramphenicol (30 µg) (Thermo Fisher Scientific, MA, USA).

### Identification of virulence factors

The virulence factors encoding genes of *P. agglomerans* KM1 were investigated using the BLASTn method against the virulence factor database (VFDB) [50] with an identity cut-off of 80%. For virulence gene identification against the VFDB, an E-value <1.0 E^-6^ was set for BLAST searches. The identified hallmarks of type VI secretion system effectors, the *Hcp* and *VgrG* genes, were validated using polymerase chain reaction with the following primers: Hcp-F: TGTAAACCAGCGCCATCAGT; Hcp-R: ACCGGTAATGCACAGCTGAA; VgrG-F: TGAATCCGCTTGCTTCCTGT; VgrG-R: ATATCGCCCATGCGTTCCAT. The primers used in the study were designed using NCBI Primer-BLAST. The thermal cycling conditions consisted of an initial denaturation at 95°C for 5 min, followed by 25 cycles of denaturation at 95°C for 30 sec, annealing at 55°C for 30 sec, and elongation at 72°C for 2 min, with a final extension at 72°C for 10 min. The PCR products were gel purified and sequenced using an ABI3730XL machine (Applied Biosystems, USA) at Cosmo Genetech (Seoul, South Korea). The sequences were identified using BLASTx program.

### Immuno-stimulations and inhibition studies

The heat-inactivated stock of *P. agglomerans* KM1 was prepared by growing bacterial culture in LB medium to an optical density of 0.9 measured at 600 nm. Bacterial cells were spun down for 5 min and re-suspended in 1 ml of a sterile and endotoxin free Dulbecco’s PBS (Welgene, Gyeongsangbuk-do, South Korea). Freshly prepared bacterial cells at concentration of 10^8^/ml were heat inactivated for 3 hours at 65°C and subsequently used to stimulate murine macrophage RAW 264.7 cell line (ATCC TIB 71, American Type Culture Collection, VA, USA) *in vitro*. The RAW 264.7 macrophages were grown in High glucose Dulbecco’s Modified Eagle Medium (DMEM, Capricorn Scientific, Germany) supplemented with 10% heat-inactivated fetal bovine serum (Atlas Biologicals, CO, USA), 100 U.mL-1 Penicillin, and 100 µg.mL-1 Streptomycin (Capricorn Scientific, Germany). 10^6^ RAW 264.7 cells were stimulated with 10^6^ heat-inactivated *P. agglomerans* or 1 ug/ml of the *E. coli* LPS-EK Ultrapure (Invivogen, CA, USA). This commercial LPS preparation was shown to activate only the TLR4 signaling. All stimulations were performed in a final volume of 1 ml. The stimulated cells were incubated for 24 hr at 37°C, afterwards culture supernatants were collected and stored at -20°C for cytokine ELISA. For inhibition studies of immunostimulatory properties of *P. agglomerans* different inhibitors of intracellular pathways were used. The involvement of the NF-κB was tested using irreversible inhibitor Bay 11-7082 at 10 µM (Invivogen, CA, USA). The action of MAPK Kinases (MEK1 and MEK2) was blocked by the use of a selective inhibitor UO126 at 5 µM (Invivogen, CA, USA). The involvement of the TLR1/2 in signal transduction was tested using CU-CPT22 at 10 µM (Tocris Bioscience, UK). The used concentrations of inhibitors were selected based on recommendations provided by the manufacturers. The TLR4 knock-out RAW 264.7 cell line (RAW-Dual™ KO-TLR4, Inivogen, CA, USA) was used to evaluate the involvement of the TLR4 signaling pathway as these cells do not respond to TLR4 agonists. The inhibition studies were conducted using 10^6^ RAW 264.7 cells or TLR4 knock-out RAW 264.7 cell line that were first seeded for 30 min in 24 well plates followed by 30 min pre-incubation with various inhibitors. Next, 10^6^ heat-inactivated *P. agglomerans* bacterial cells were added to pre-treated RAW 264.7 cells in a final volume of 1 ml. The culture supernatants collected from non-stimulated cells and cells that were treated only with inhibitors were used as negative controls. The cultures were incubated for 24 hours, and afterwards supernatants were collected and used for cytokine quantification in ELISA.

### Quantification of cytokines by ELISA and nitrite detection

The concentrations of TNF-α, IL-6, and IL-10 in culture supernatants were measured using ELISA Max Deluxe were from Biolegend^®^ (CA, USA) according to manufacturer’s instructions. Taking into account various detection limits of used ELISAs, culture supernatants were diluted 50 times for TNF-α, 100 times for IL-6, and 2 times for detection of IL-10 secretion. The detection limits for different cytokines were as follows (TNF-α/IL-6: 500 pg/ml, IL-10: 2.000 pg/ml). The nitrite concentrations were detected in culture supernatants using Griess reaction and a commercial kit from Promega (WI, USA) according to manufacturer’s instructions. In this assay, undiluted culture supernatants were used after 24 hrs of stimulation of RAW 264.7 cells or TLR4 knock-out RAW 264.7 cells with *P. agglomerans* whole-cell preparation in the presence or absence of various inhibitors.

### Statistical analysis

All values recorded in experiments were presented as mean ± standard error of the mean (SEM) of three independent experiments. Statistical analysis was conducted using a Student’s t-test, with GraphPad Prism 5.0c software. Values with P < 0.001 were considered as significantly different.

## Results

### Genomic features of *Pantoea agglomerans* KM1 isolate

Whole-genome sequencing of *P. agglomerans* KM1 using the Illumina HiSeq 4000 platform resulted in approximately 168-fold genome coverage. Collectively, 6,890,002 raw reads of 101 bp in length were subjected to quality filtering with low quality scores (≤Q27) and a minimum length of 36 bases. After trimming, 6,323,946 (91.78%) clean reads were used for downstream analysis. *De novo* assembly of the sequence reads resulted in 46 contigs, and the N50 of the contigs was 330,495 bp. The maximum length of a contig was 1,299,234. BUSCO provides quantitative measures for the assessment of genome assembly based on evolutionarily informed expectations of gene content from near-universal single-copy orthologs. Evaluation of a genome completeness using BUSCO revealed that *P. agglomerans* KM1 was 99.2% complete, suggesting that most of the recovered genes could be classified as complete and single-copy. Moreover, the genome contained one fragmented BUSCO contributing to 0.8% of the total BUSCO groups (S1 Fig). Hence, the draft genome sequence of *P. agglomerans* KM1 included one circular DNA chromosome of 4,039,945 bp with a GC content of 55.55%. The KM1 chromosome contained 4,651 genes, of which 4488 were protein coding, 64 tRNAs, 1 rRNA and 9 ncRNAs. In addition to the bacterial chromosome, the genome consisted of three mega plasmids named pKM1_1, pKM1_2 and pKM1_3 of sizes 562,844 bp, 222,325 bp and 170,642bp, respectively. The number of protein-coding genes present on pKM1_1, pKM1_2, and pKM1_3 was 538, 237 and 152, respectively.

### Genome annotation

The predicted Open Reading Frames (ORFs) were further classified into COG functional groups and SEED Subsystem categories. The top 5 most abundant COG category based on counts are class of R (702 ORFs, General function prediction only), class of E (270 ORFs, Amino acid transport and metabolism), class of C (258 ORFs, Energy production and conversion), class of J (245 ORFs, Translation, ribosomal structure and biogenesis) and class of L (238 ORFs, Replication, recombination and repair) categories. For SEED subsystem analysis, 36,697 ORFs were classified to SEED Subsystem categories (S3 Fig). Among the SEED categories, the top five most abundant categories were Carbohydrates (5865 ORFs), Amino Acids and Derivatives (5115 ORFs), Cofactors, Vitamins, Prosthetic Groups, Pigments (3026 ORFs), Protein Metabolism (2490 ORFs) and DNA metabolism (2242 ORFs) (S2 Fig). Fig 1 shows the circular genome maps of the KM1strain showing all annotated ORFs and GC content using the GView program.

**Fig 1.**
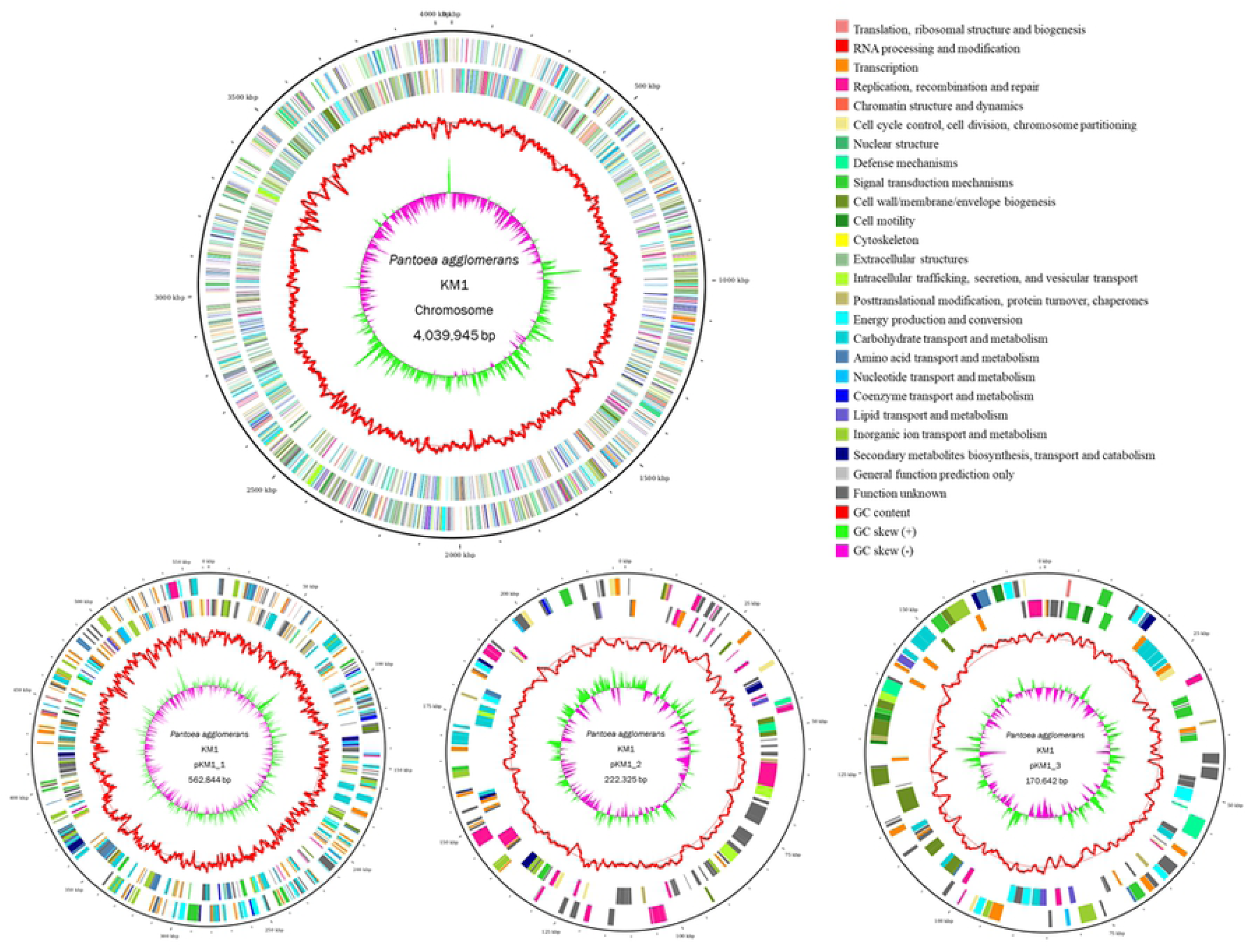
The circular genome maps of the *P. agglomerans* KM1 draft genome. Genome map showing the features of *P. agglomerans* KM1 chromosome and plasmids pKM1_1, pKM1_2, and pKM1_3. Circles illustrate the following from outermost to innermost rings: (1) forward CDS, (2) reverse CDS, (3) GC content, and (4) GC skew. All the annotated ORFs are colored differently according to the COG assignments.

### Phylogenetic analysis and comparative genomics

The degree of genomic similarity of *P. agglomerans* KM1 with closely related *Pantoea* species was estimated using the OrthoANI tool. Analysis of the orthologous average nucleotide identity (OrthoANI) values among *Pantoea* genome sequences revealed that strain KM1 had 79.12% to 98.66% genome sequence similarities with other *Pantoea* species (Fig 2). The KM1 genome was most closely related to that of *P. agglomerans* UAEU18 (98.68%), a plant-growth promoting bacterium isolated from date palm rhizosphere soil in the United Arab Emirates [51]. This was followed by *P. agglomerans* C410P1 (98.66%), a biocontrol agent isolated from lettuce in China (GenBank accession number CP016889), *P. agglomerans* Tx10 (97.44%), isolated from the sputum of cystic fibrosis patient [52], *P. agglomerans* JM1 (97.41%), isolated from soil polluted by beta-lactam antibiotics [53], *P. agglomerans* IG1 isolate (97.39%), an immunopotentiator producer [54] and *P. agglomerans* P5 (97.38%), a plant growth-promoting rhizobacterium [17]. Furthermore, the KM1 strain is 90.80% similar to that of *Pantoea vagans* C9-1, which is a biocontrol agent used against fire blight of apple and pear trees [55], and 90.92% similar to that of *Pantoea vagans* MP7, which was isolated from fungus-growing termites in South Africa [9]. Therefore, comparative whole-genome sequence analysis indicated that KM1 strain belongs to the species *Pantoea agglomerans* with a species cut-off level of >97% [56]. Phylogenetic tree analysis based on 16S rRNA gene sequences revealed that KM1 clustered closely with *P. agglomerans* C410P1 and UAEU18 (S2 Fig). This coincides with results obtained using the phylogenetic tree relationship analysis based on OrthoANI values with 98.66% and 98.68% similarity, respectively, suggesting a common evolutionary origin.

**Fig 2.**
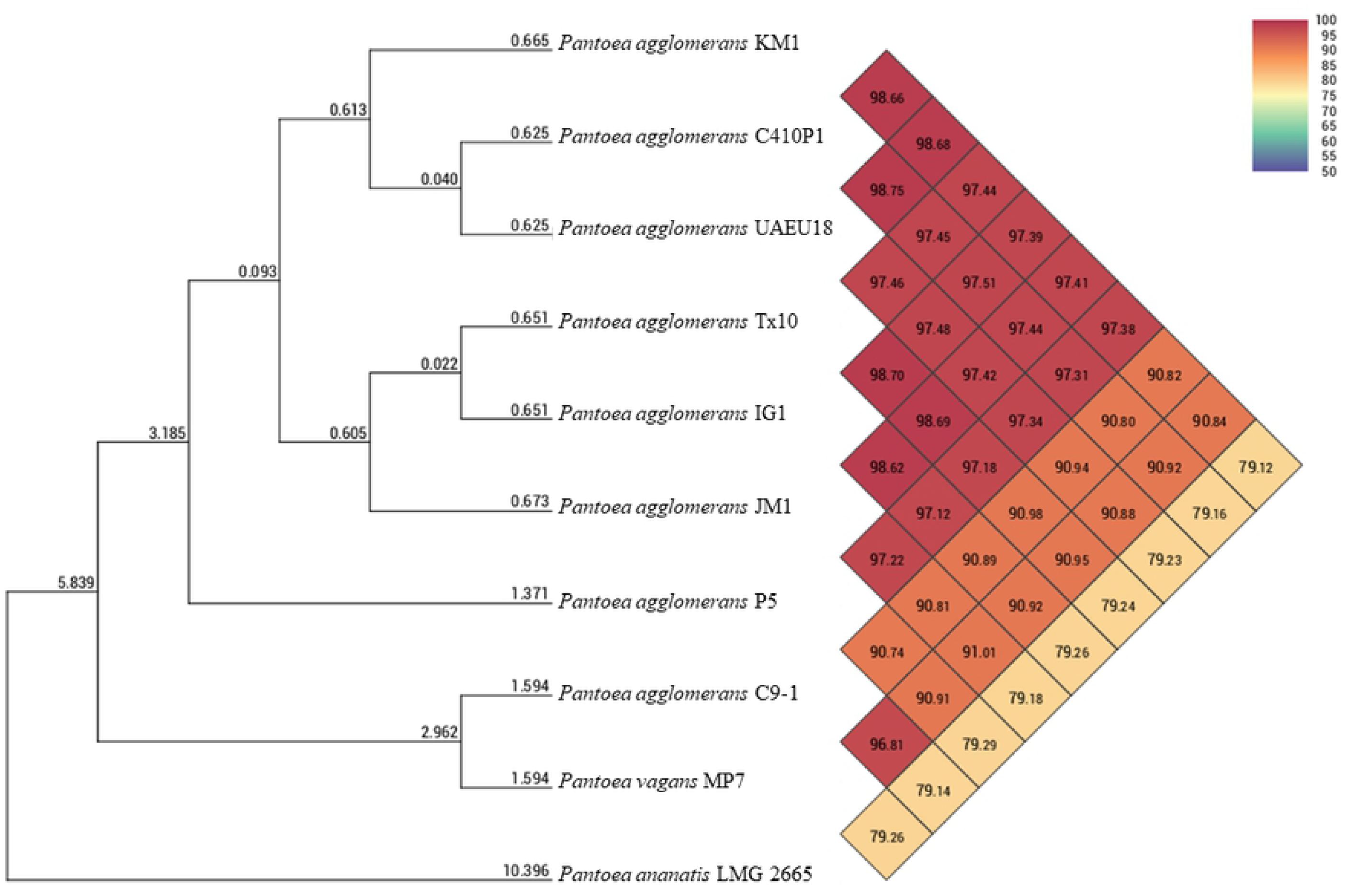
Phylogenetic analysis of *P. agglomerans* KM1 based on average nucleotide identity. Heatmap showing the orthologous average nucleotide identity (OrthoANI) values between *P. agglomerans* KM1 genome and its closely related species. The OrthoANI values between KM1 and other *Pantoea* genomes were as follows: *P. agglomerans* C410P1 (98.66%), *P. agglomerans* UAEU18 (98.68%), *P. agglomerans* Tx10 (97.44%), *P. agglomerans* IG1 (97.39%), *P. agglomerans* JM1 (97.41%), *P. agglomerans* P5 (97.38%), *P. vagans* C9-1 (90.82%), *P. vagans* MP7 (90.84%) and *P. ananatis* LMG 2655 (79.12%). Values greater than 97% indicate that strains belong to the same species.

Comparative genomic analysis was performed to extend our understanding of the genomic characteristics of *P. agglomerans* KM1. The genome synteny between the KM1 genome and its closely related *P. agglomerans* strains C410P1, UAEU18, TH81 and L15 is shown in Fig 3. The KM1 chromosome and contigs associated to three-mega plasmids show high collinearity with that of C410P1. The genome alignment revealed the presence of 36 collinear blocks and several regions of translocations and inversions. In addition, the chromosomal alignments between strains C410P1 and KM1 are nearly identical, as shown by the presence of large collinear blocks of high similarity when most portions of the two chromosomes are aligned onto each other. Notably, blocks seven and eight of KM1 display translocations, suggesting a different syntenic relationship to the reference genome sequence. Interestingly, the three plasmids of KM1 are syntenic with the three plasmids of C410P1 strain, where in plasmid pKM1_3 shows inversions relative to C410P1.

**Fig 3.**
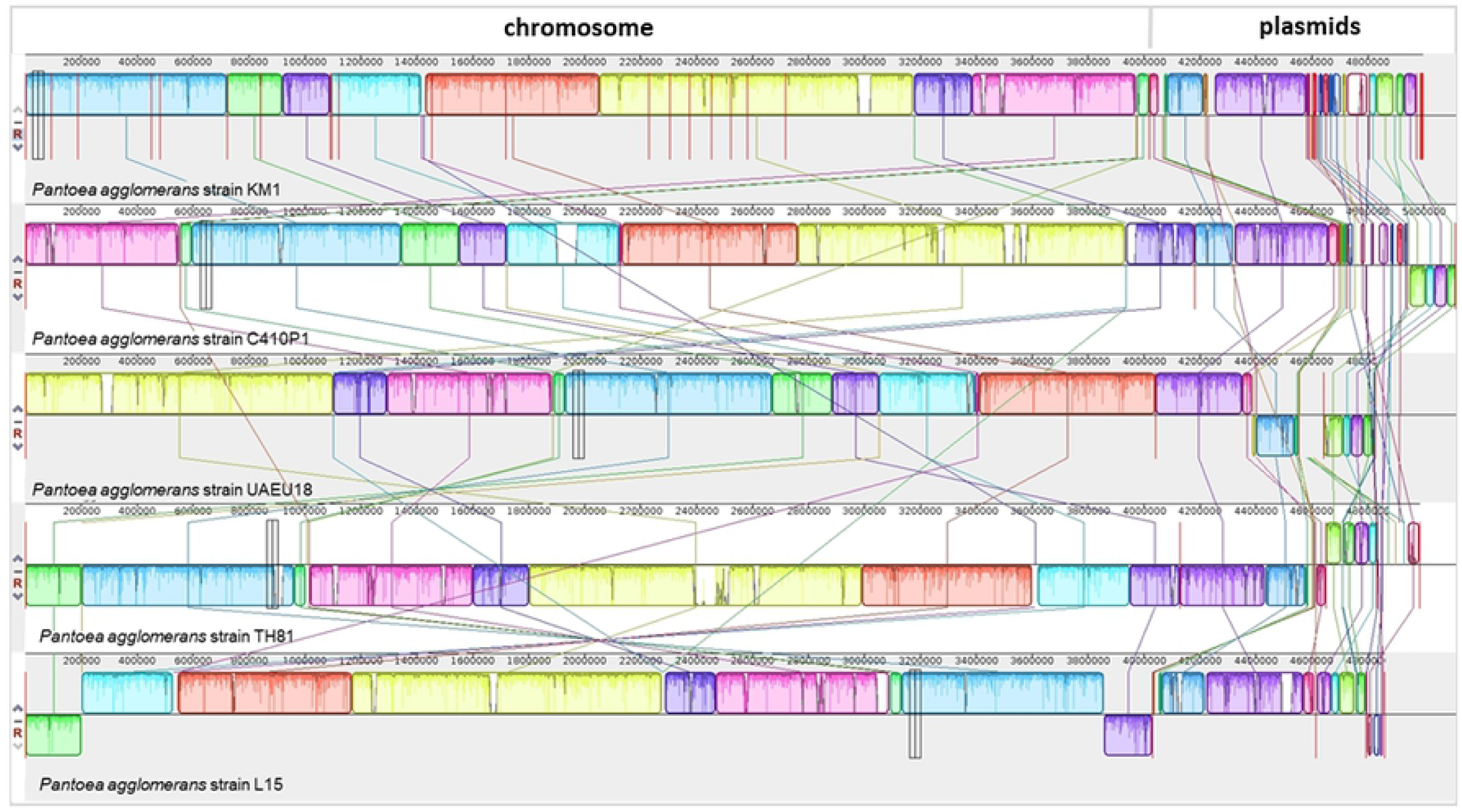
Genome alignments showing synteny blocks among *P. agglomerans* strains obtained using progressive Mauve. *P. agglomerans* KM1 were compared with other closely related strains namely C410P1, UAEU18, TH81 and L15. Each genome is laid out horizontally with homologous segment outlined as colored rectangles. Each same color block represents a locally collinear block (LCB) or homologous region shared among genomes. Rearrangement of genomic regions was observed between the two genomes in terms of collinearity. Inverted regions relative to KM1 are localized in the negative strand indicated by genomic position below the black horizontal centerline in the Mauve alignment.

A pan-genome map for the KM1 strain and four other completely sequenced *P. agglomerans* strains was drawn using PGAP and Gview server (Fig 4). Based on Roary-matrix pipeline, the pan-genome analysis of five *P. agglomerans* strains revealed 6,727 protein coding genes comprising the pan-genome, of which 3,065 genes (45.6%) corresponded to the accessory genome and 3,662 genes (54.4%) corresponded to the core genome (S4 Fig). The core genome from each species was used to construct the phylogenetic tree. Roary matrix-based phylogenetic tree analysis revealed that KM1 is closely related to C410P1. As the *P. agglomerans* pan-genome increases with the addition of new strains, the size of core genome decreases, suggesting that *P. agglomerans* has an open pan-genome [57] (S5 Fig).

**Fig 4.**
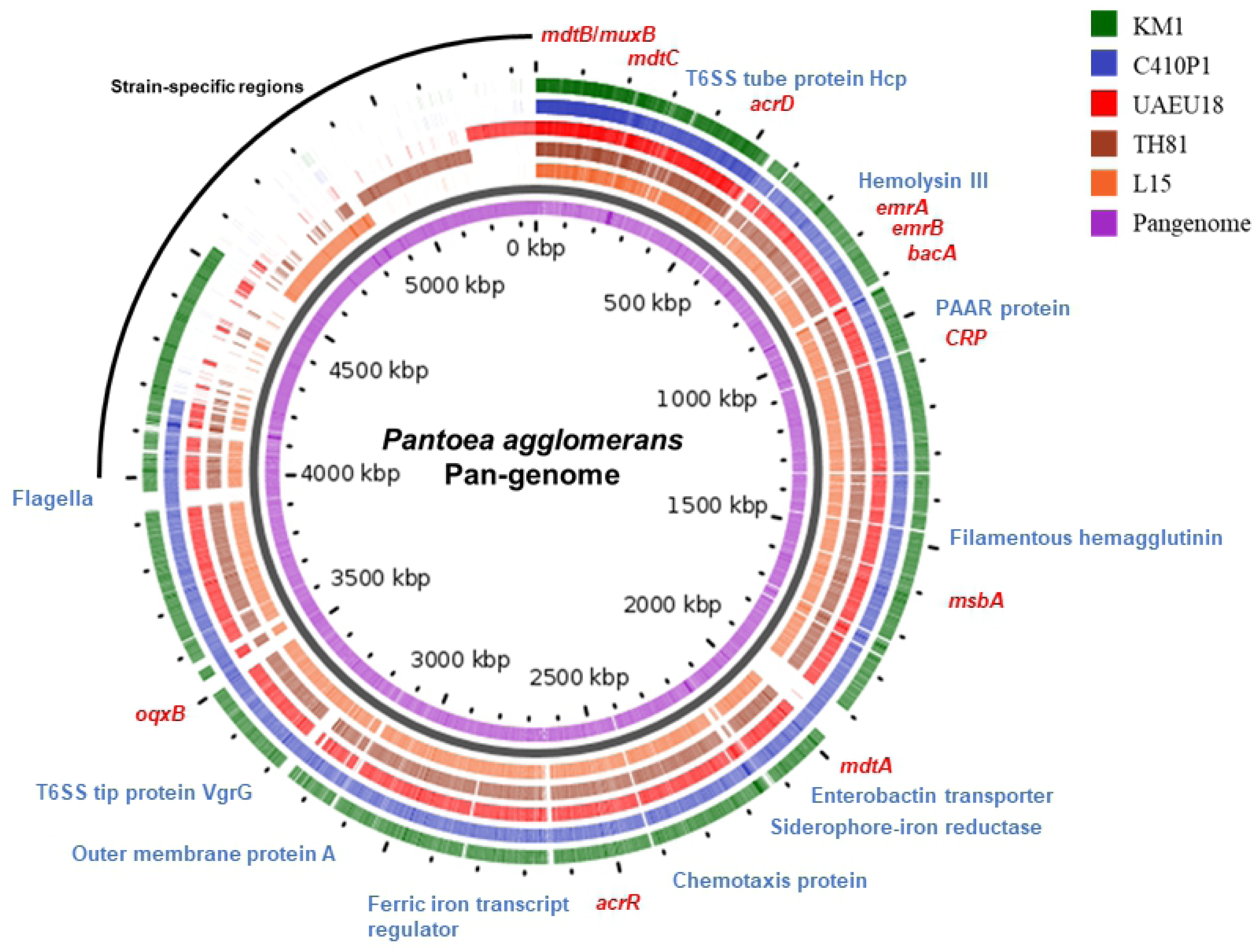
Pan-genome analysis of *P. agglomerans* strains obtained using Gview server. The innermost circle shows the pan-genome (purple), and outer circlers indicate the genomes of *Pantoea agglomerans* strains L15 (orange), TH81 (brown), UAEU18 (red), C410P1 (blue), and KM1 (green). Genes with specialized functions were labelled with different colors: virulence-related genes (blue), antibiotic resistance genes (red), and strain-specific regions (black).

### Genomic sequences involved in response to viruses and horizontal gene transfer

Using PHASTER, two prophage regions were predicted in KM1 chromosome, where genomic island (GI) regions 6 to 13 coincide with prophage region 1, and GI region 20 to 21 incorporates prophage region 2 (Fig 5 and S6 Fig). Prophage region 1 is an intact prophage with a size of 52 kb, a GC content of 52.56%, and 82 ORFs in the phage protein database. Prophage region 2 is an incomplete prophage and it has a size of 12.9 kb, with a GC content of 47.26%, and 13 ORFs. Several genes that have a phage-derived genetic material were present in the KM1 chromosome such as a phage integrase, a phage lysozyme, a phage terminase, a phage tail tip, a phage capsid and a scaffold protein. The *P. agglomerans* KM1 genome also contained six CRISPR loci (CRISPR1-CRISPR6) with direct repeats ranging between 23 to 42 and mean size spacers ranging from 35 to 59 (S1 Table). Additionally, 22 GIs were identified on the chromosome of *P. agglomerans* KM1 (Fig 5 and S2 Table), suggesting horizontal DNA transfer. In the KM1 chromosome, GI region 4 was found to encode type I secretion proteins and GI region 16 encodes LysR-family transcriptional regulator, which are required for virulence. GI region 1 contained glutathione S-transferase, omega (EC 2.5.1.18), which serve as the primary defenses against oxidative stress [58] (S2 Table). Additionally, 4, 2, and 2 GIs were detected on KM1 plasmids pKM1_1, pKM1_2, and pKM1_3, respectively (Fig 5). Plasmid pKM1_1 harbored genes involved in carotenoid biosynthesis including lycopene cyclase (*CrtY*), zeaxanthin glucosyltransferase (*CrtX*) and geranylgeranyl diphosphate synthase (*CrtE*), suggesting that *P. agglomerans* KM1 is a carotenogenic bacterium, where carotenoids serve an essential role in protection against oxidative stress and protection from oxygen during nitrogen fixation [59] (S2 Table).

**Fig 5.**
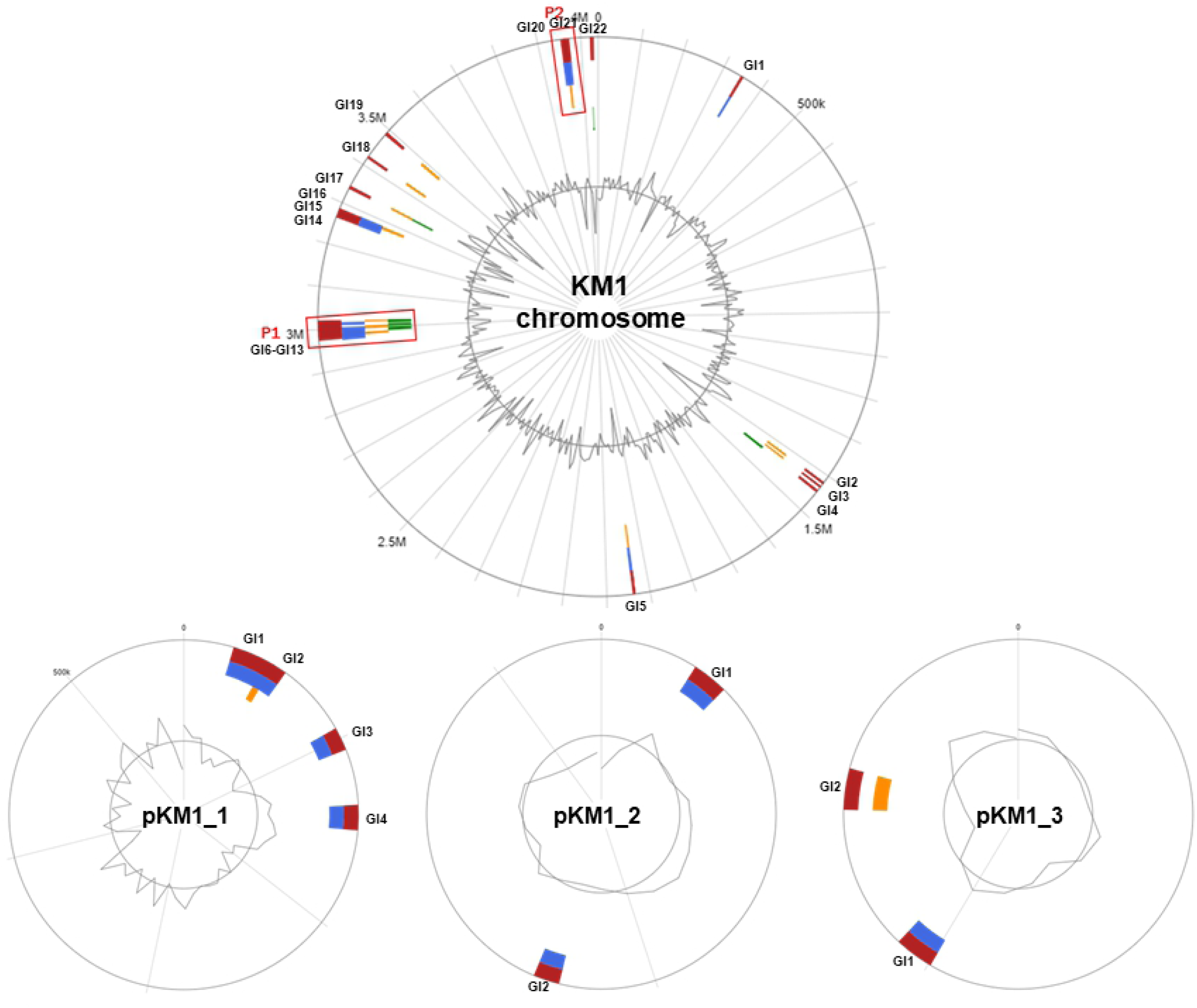
Genomic islands (GIs) in *P. agglomerans* strain KM1 predicted using IslandViewer4. The predicted genomic islands are colored based on the prediction methods. Red indicates an integrated analysis, blue represents IslandPath-DIMOB prediction, orange represents SIGI-HMM prediction, and green indicates IslandPick analysis. The circular plots show the genomic islands in *P. agglomerans* KM1 chromosome, and plasmids pKM1_1, pKM1_2 and pKM1_3. GIs are labelled in blue. Prophage regions predicted by PHASTER were indicated in red boxes for prophage region 1 (P1) and region 2 (P2).

### Biochemical characterization

Biochemical analysis of the *P. agglomerans* KM1 was conducted using an API 20E test (S3 Table). The isolate was catalase positive and scored negative in the oxidase and urease test. KM1 did not decarboxylate lysine and ornithine, nor produced hydrogen sulfide (H_2_S) and indole. In addition, the isolate was able to ferment D-glucose, D-mannitol, L-rhamnose, D-sucrose, Amygdalin, L-arabinose and Lactose. The API 20E test was negative for arginine dihydrolase, ornithine decarboxylase, lysine decarboxylase, and tryptophan deaminase. The KM1 isolate had no ability to ferment inositol, D-sorbitol, and D-melibiose, however, it showed positivity in acetoin production, β-galactosidase test, citrate utilization test, and possessed gelatinase activity. Obtained numerical profile confirmed that isolated KM1 strain belongs to *P. agglomerans* species.

### Genes involved in antibiotic resistance

The presence of Antibiotic Resistance Genes (ARGs) was identified using BLAST against the CARD reference sequences (S4 Table). We identified 12 ARGs in KM1 chromosome and 1 ARG in plasmid pKM1_3 showing greater than 70% identity to well-characterized ARGs in CARD database. The ARGs identified in KM1 were classified into five gene families including resistance-nodulation-cell division (RND) antibiotic efflux pump (*CRP, oqxB, mdtA, mdtB, mdtC, acrR, acrD*, and *MuxB*), ATP-binding cassette (ABC) antibiotic efflux pump (*msbA*), major facilitator superfamily (MFS) antibiotic efflux pump (*emrA* and *emrB*), undecaprenyl-pyrophosphate related protein (*bacA*) and family of phosphoethanolamanine transferase (*arnA*). These genes provide resistance to multiple antibiotic classes such as macrolides, fluoroquinolones, tetracyclines, aminoglycosides, nitroimidazoles aminocoumarins, and peptide based antibiotics. The predicted multidrug resistant genes present in KM1 genome were primarily involved into two resistance mechanisms namely antibiotic efflux and antibiotic target alteration. These results are consistent with data obtained from the Kirby-Bauer disk diffusion test showing that KM1 strain is resistant to penicillin G, vancomycin, bacitracin, fosfomycin, and rifampicin (Table 1).

**Table 1.**
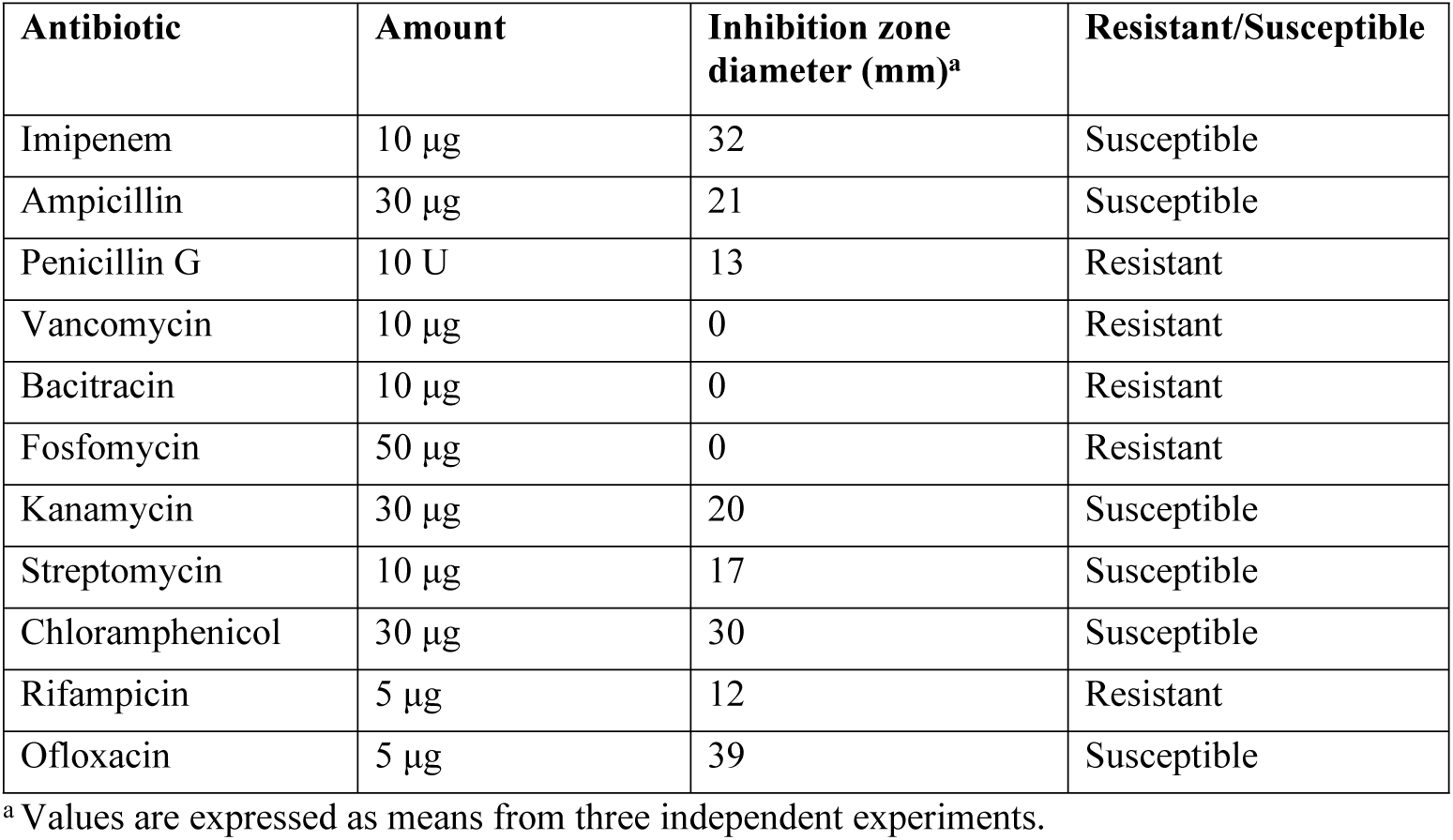
Antibiotic susceptibility profile of *P. agglomerans* KM1.

### Genes involved in adaptations to environmental stress

In the past, *P. agglomerans* strains have been isolated from a multitude of environments such as extreme desiccation in powdered infant milk formula [10], broad range of temperature (3 – 42 °C) or pH (5-8.6) regimes [60] and osmotically challenging environments [61], as such its survival in stressful environments have not been examined so far at the molecular level. Table 2 demonstrates the list of genes found in the KM1 isolate, associated with resistance to stressful environmental conditions. Several genes encode products involved in osmoregulation included aquaporin Z (*aqpZ*), osmotically inducible protein (*OsmY*), potassium transporter protein (*TrkH* and *TrkA*), glycine betaine transporter protein (*proP*), trehalose-6-phosphate synthase (*otsA* and *otsB*), and glutathione-regulated potassium-eflux system protein (*kefB*). Further in-depth analysis revealed the presence of genes coding for proteins involved in adaptation to temperature fluctuations such as cold shock proteins (*cspA, cspE*, and *cspD*) and heat shock proteins (*DnaJ, DnaK, GrPE, hslR, IbpA, hspQ*). Finally, genes conferring resistance to oxidative stress were also detected in KM1 strain such as catalase (*katE, katG*), superoxide dismutase (*sodA*), glutathione S-transferase (*GST*), glutathione peroxidase (*GPX*), and DNA protection during starvation protein (*Dps*).

**Table 2.** Genes associated with resistance to environmental stress in *P. agglomerans* KM1.

| Environmental stress resistance | Annotation | Locus tag |
| --- | --- | --- |
| Osmotic stress |  |  |
| *OsmY* | Osmotically-inducible protein Y | HBB05_RS01050 |
| *aqpZ* | Aquaporin Z | HBB05_RS05300 |
| *TrkH*, *TrkA* | Trk system potassium transporter protein | HBB05_RS05080,<br>HBB05_RS02110 |
| *proP* | Glycine betaine/L-proline transporter | HBB05_RS19460,<br>HBB05_RS02985 |
| *proQ* | RNA chaperone ProQ | HBB05_RS14815 |
| *otsA*, *otsB* | Trehalose-6-phosphate synthase | HBB05_RS15190,<br>HBB05_RS15195 |
| *kefB*, | Glutathione-regulated potassium-efflux system protein | HBB05_RS02355,<br>HBB05_RS02360 |
| *TreF* | Cytoplasmic trehalase | HBB05_RS00975 |
| Heat shock |  |  |
| *DnaJ*, *DnaK* | Molecular chaperone DnaJ | HBB05_RS18860,<br>HBB05_RS19155,<br>HBB05_RS07025,<br>HBB05_RS07020 |
| *hslR* | Ribosome-associated heat shock protein Hsp15 | HBB05_RS02565 |
| *IbpA*, *hspQ* | Heat shock protein | HBB05_RS04290,<br>HBB05_RS10600 |
| *GrpE* | Nucleotide exchange factor | HBB05_RS18265 |
| Cold shock |  |  |
| *cspA*, *cspE*,<br>*cspD* | Cold shock protein | HBB05_RS19980,<br>HBB05_RS03250,<br>HBB05_RS09100,<br>HBB05_RS10245 |
| Oxidative stress |  |  |
| *dps* | DNA protection during starvation protein | HBB05_RS09995,<br>HBB05_RS00870 |
| *oxyR*, | DNA-binding transcriptional regulator | HBB05_RS03440 |
| *Gpx* | Glutathione peroxidase | HBB05_RS19175,<br>HBB05_RS03445,<br>HBB05_RS12185 |
| *Gst* | Glutathione S-transferase | HBB05_RS18795,<br>HBB05_RS18945,<br>HBB05_RS19010,<br>HBB05_RS20560,<br>HBB05_RS01550,<br>HBB05_RS03195,<br>HBB05_RS03320,<br>HBB05_RS04480,<br>HBB05_RS05500 |
| *KatE*, *katG* | Catalase | HBB05_RS12490,<br>HBB05_RS16130 |
| *sodA* | Superoxide dismutase | HBB05_RS04520,<br>HBB05_RS12380 |

### Genome-based identification of virulence factors

The putative virulence factors in *P. agglomerans* KM1 were predicted by BLAST searches against the VFDB and genome annotation using NCBI PGAP. As demonstrated in Table 3, identified virulence factors were classified into five categories including secretion systems, adhesion, motility, iron uptake/sequestration system and toxins.

**Table 3.** Virulence factors of *P. agglomerans* KM1.

| Virulence factor | Annotation | Locus tag |
| --- | --- | --- |
| **Secretion system** |  |  |
| *TssA, TssB, TssC, TssD (Hcp), TssE, TssF, TssG, TssH (ClpV), TssI (vgrG), TssJ, TssK, TssL (DotU), TssM (IcmF), tagF, tagH, PAAR* | Type VI secretion protein | HBB05_RS11570, HBB05_RS11575, HBB05_RS11580, HBB05_RS11585, HBB05_RS11590, HBB05_RS11595, HBB05_RS11600, HBB05_RS11605, HBB05_RS11615, HBB05_RS11650, HBB05_RS11670, HBB05_RS11675, HBB05_RS11680, HBB05_RS11685, HBB05_RS11700, HBB05_RS11705, HBB05_RS11710, HBB05_RS11900, HBB05_RS11905, HBB05_RS11910, HBB05_RS17335, HBB05_RS17360, HBB05_RS17365, HBB05_RS17370, HBB05_RS17375, HBB05_RS17380, HBB05_RS17385, HBB05_RS17410, HBB05_RS17415, HBB05_RS17420, HBB05_RS17435, HBB05_RS17440, HBB05_RS17445, HBB05_RS18610, HBB05_RS18655, HBB05_RS17395 |
| **Adhesion** |  |  |
| *fha* | Filamentous hemagglutinin | HBB05_RS03515, HBB05_RS04420, HBB05_RS11225, HBB05_RS15615 |
| *ompA* | Outer membrane protein A | HBB05_RS19865, HBB05_RS04215, HBB05_RS10555, HBB05_RS11915, HBB05_RS17425 |
| **Motility** |  |  |
| *fliC, fliD, fliE, fliF, fliG, fliH, fliI, fliJ, fliK, fliL, fliM, fliN, fliO, fliQ, fliS, fliT, fliZ, flhA, flhC, flhD, flhE, flgA, flgB, flgC, flgD, flgE, flgF, flgG, flgH, flgI, flgJ, flgK, flgL, flgN, motB* | Flagella | HBB05_RS15305, HBB05_RS15310, HBB05_RS15315, HBB05_RS15320, HBB05_RS15410, HBB05_RS15415, HBB05_RS15420, HBB05_RS15425, HBB05_RS15430, HBB05_RS15435, HBB05_RS15440, HBB05_RS15445, HBB05_RS15450, HBB05_RS15455, HBB05_RS15460, HBB05_RS15470, HBB05_RS15285, HBB05_RS15115, HBB05_RS15180, HBB05_RS15185, HBB05_RS15110, HBB05_RS11030, HBB05_RS11035, HBB05_RS11040, HBB05_RS11045, HBB05_RS11050, HBB05_RS11055, HBB05_RS11060, HBB05_RS11065, HBB05_RS11070, HBB05_RS11075, HBB05_RS11080, HBB05_RS11085, HBB05_RS11020, HBB05_RS15170 |
| *CheV, CheY, CheW, CheA* | Chemotaxis protein | HBB05_RS21255, HBB05_RS08110, HBB05_RS12020, HBB05_RS15130, HBB05_RS15160, HBB05_RS15165, HBB05_RS15645 |
| **Iron uptake system** |  |  |
| *fur* | Ferric iron uptake transcriptional regulator | HBB05_RS09340 |
| *EfeO* | Iron uptake system protein | HBB05_RS15240 |
| *SitC* | Iron/manganese ABC transporter permease subunit | HBB05_RS21090 |
| *fepB, fepG, entS* | Enterobactin transporter | HBB05_RS06140, HBB05_RS06155, HBB05_RS06145 |
| *fepA* | TonB-dependent siderophore receptor | HBB05_RS20475, HBB05_RS21020, HBB05_RS06180 |
| *fepD* | Ferric siderophore ABC transporter permease | HBB05_RS06150 |
| *fhuF* | Siderophore-iron reductase | HBB05_RS06780 |
| **Toxin** |  |  |
| *Hha, ShlB, FhaC, HecB, XhlA* | Hemolysin | HBB05_RS23025, HBB05_RS03510, HBB05_RS04405, HBB05_RS08630, HBB05_RS11220, HBB05_RS14060, HBB05_RS15620 |
| *hlyIII* | Hemolysin III | HBB05_RS00635 |

In the secretion system category, all the putative genes belonging to type VI secretion system (T6SS) were present. The genes encoding the complete structural gene components of the T6SS apparatus consisted of *TssA*-*TssM*, hemolysin-coregulated protein (*Hcp*), valine-glycine repeat G (*VgrG*) protein, proline-alanine-alanine-arginine repeats (*PAAR*) and *ClpV*. In addition, the presence of T6SS hallmarks *Hcp* and *VgrG* genes in KM1 was validated using PCR and amplicon sequencing (S7 Fig). The latter showed 100% sequence identity to *Hcp* and *VgrG* proteins in *P. agglomerans*. Previously, the T6SS was demonstrated to be responsible for delivery of toxic effector proteins to other microorganisms and eukaryotic cells [62, 63].

In the adhesion category, the identified genes encoded filamentous hemagglutinin (*FHA*), an attachment factor-facilitating adherence to host ciliated epithelial cells of the respiratory tract [64]. The outer membrane protein A (*OmpA*) involved in adhesion and biofilm formation [65] was also present. In the motility category, genes involved in flagella were detected in KM1, which play roles in biofilm formation, virulence factor secretion and adhesion in addition to motility [66].

In the iron uptake category, chromosomal genes encoding proteins related to iron uptake and transport included ferric iron uptake transcriptional regulator (*fur*), iron uptake protein (*EfeO*). Plasmid pKM1_3 contained the iron/manganese ABC transporter permease (*SitC*). In addition, genes related to high-affinity iron-chelating molecules i.e. siderophores, encompassed the enterobactin transporter (*fepB, fepG, entS*), TonB-dependent siderophore receptor (*fepA*), ferric siderophore ABC transporter permease (*fepD*, and siderophore-iron reductase (*fhuF*). These molecules are crucial in sequestrating iron from the host during infection [67].

The last group of virulence factors contained genes coding for extracellular cytotoxic proteins hemolysin III, hemolysin *XhlA*, and hemolysin secretion/activation genes (*Hha, ShlB, FhaC, HecB*). The cytotoxic activity of these molecules involves binding to erythrocyte surface followed by a pore formation and cell lysis. The mentioned above bacterial secretion system I was shown to be involved in extracellular transport of bacterial toxins such as hemolysins.

### Immuno-stimulatory potential of *P. agglomerans KM1*

In order to supplement the genomic virulence study with an additional data showing potential contribution of *P. agglomerans* KM1 to induction of inflammation, the immuno-stimulatory properties of the KM1 isolate were tested *in vitro* using the RAW 264.7 macrophage cell line. The latter was stimulated with a heat-inactivated whole-cell preparation of the KM1 isolate followed by the measurement of cytokine secretion and Nitric Oxide (NO) production. The mechanisms underlying KM1-associated production of cytokines and NO were tested by inhibiting TLR1/2 and TLR4 signaling and activation of NF-kB and MAPK Kinases, such as MEK 1 and MEK2.

As shown in Fig 6A, stimulations of RAW 264.7 macrophages with *P. agglomerans* KM1 resulted in the production of TNF-α. Under identical stimulation conditions RAW 264.7 TLR4 knock-out cells exhibited a significantly reduced TNF-α secretion. Analysis of a possible involvement of the TLR1/2 signaling in TNF-α secretion by stimulated RAW 264.7 macrophages showed partial reduction in the presence of the CU-CPT22 TLR-1/2 inhibitor. However, this inhibitory effect was itself significantly less potent than the effect observed in the TLR4 knock-out condition. Adding CU-CPT22 to stimulated TLR4 knock-out cells did not result in an additional inhibitory effect, indicating a dominant role of the TLR4 signaling. The use of inhibitors of intracellular pathways leading to activation of the NF-κB (Bay 11-7082) and MAPKKs (UO126) allowed to further analyze mechanisms underlying KM1 associated TNF-α secretion. The use of Bay 11-7082 resulted in a significant reduction of TNF-α secretion by RAW 264.7 cells, similar to the inhibition observed in the absence of the TLR4 receptor. There was no added effect of the inhibitor in the TLR4 knock-out stimulation setting. Finally, we did not observe any inhibitory effect of the UO126, indicating a lack of the involvement of MAPK Kinases in *P. agglomerans* induced TNF-α secretion. In order to assess the results, the experimental setting was accompanied by a series of control conditions. First, RAW 264.7 cells and the RAW 264.7 TLR-4 knock-out cell line were both stimulated with ultra-pure LPS, showing the effectiveness of the absence of TLR4 in the inhibition of TNF-α secretion (Fig 6B). Under non-stimulated conditions, neither of the cell lines produced any measurable level of TNF-α. Finally, none of the inhibitors used in this study was able to drive TNF-α secretion by itself (Fig 6C)

**Fig 6.**
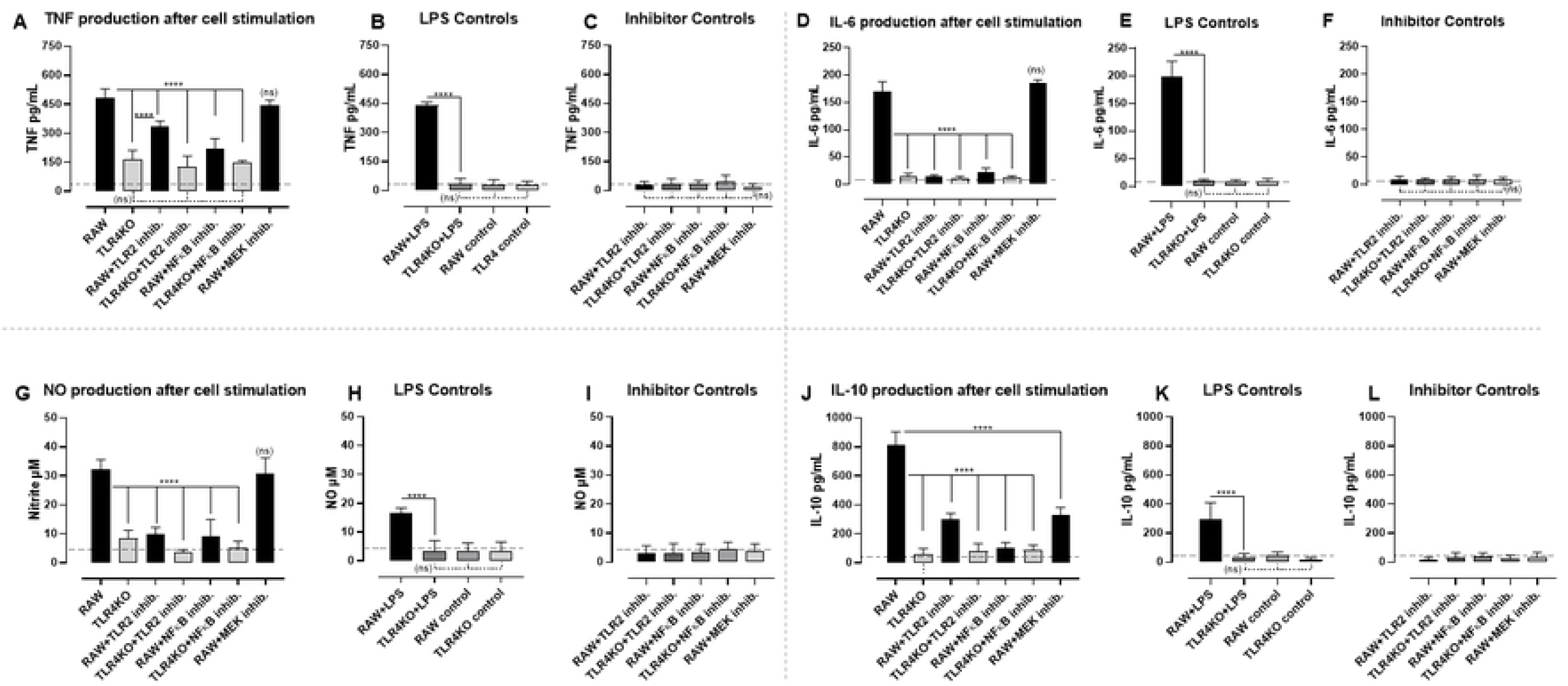
Cytokine and nitrite production by RAW 264.7 macrophages stimulated with a heat inactivated whole-cell *P. agglomerans*. Panels A, D, G and J show TNF-α, IL-6, Nitrite and IL-10 secretion by the simulated RAW 264.7 cells (RAW), TLR4 knock-out RAW 264.7 cells (TLR4KO), RAW 264.7 cells (RAW) in combination with TLR2 inhibitor (RAW+ TLR2 inhib), TLR4 knock-out RAW 264.7 cells in combination with TLR2 inhibitor (TLR4KO + TLR2 inhib.). Additionally stimulations were performed using RAW 264.7 cells in combination with NFκB inhibitor (RAW+NFκB inhib.), TLR4 knock-out RAW 264.7 cells in combination with NFκB inhibitor (TLR4KO + NFκB inhib.) and RAW 264.7 cells in combination with MEK1/2 inhibitor (RAW + MEK inhib.). Panels B, E and K, include control conditions: RAW 264.7 cells stimulated with TLR4 agonist ultrapure LPS (RAW + LPS), TLR4 knock-out RAW 264.7 cells stimulated with TLR4 agonist ultrapure LPS (TLR4KO + LPS), non-stimulated RAW 264.7 cells (RAW control), non-stimulated TLR4 knock-out RAW 264.7 cells (TLR4 control). Panels C, F, I and L include control stimulations: RAW 264.7 cells and TLR4 knock-out RAW 264.7 cells in the presence of inhibitors alone, RAW + TLR2 inhibitor, TLR4KO + TLR2 inhibitor, RAW + NFκB inhibitor, TLR4KO + NFκB inhibitor, and RAW + MEK inhibitor. Values with P < 0.001 were considered as significantly different (****).

In order to evaluate the capacity *P. agglomerans* to trigger IL-6 production by RAW 264.7 cells, an identical experimental set-up was used as the one described above. Here, the results presented in Fig 6D show that high levels of cytokine production were obtained by *P. agglomerans* stimulation of RAW 264.7 cells, but not RAW 264.7 TLR4 knock-out cells. In contrast to the partial inhibition of TNF-α production by CU-CPT22, addition of this TLR1/2 inhibitor resulted in a complete abolishment of IL-6 production. The same level of inhibition was also obtained by blocking the NF-κB signaling pathway alone or in combination with inhibition of TLR function. Blocking of MAPK Kinases did not result in reduction of IL-6 production. As for the TNF-α induction experiment, all necessary control culture conditions were included, showing the potency of the TLR4 knock-out construct in the inhibition of LPS-induced cytokine production (Fig 6E), as well as the lack of significant IL-6 induction by any of the inhibitors alone, used in this study (Fig 6F).

Next, the secretion of NO, an important pro-inflammatory effector molecule was analyzed using the same experimental setting. The presence of NO in culture supernatant can be measured using Griess reaction recording levels of a nitrite. As showed in Fig 6G, while *P. agglomerans* stimulated RAW 264.7 cells produced NO, interrupting either TLR4 or TLR1/2 signaling, resulted in a significant reduction of NO production. Also here, inhibition of NO production was obtained by blocking of the NF-κB signaling, but not the MAPK Kinase pathway. As for the experiments outlined above, all cell culture control conditions were included, allowing proper result interpretation (Fig 6H and Fig 6I.)

Finally, the potential of *P. agglomerans* to induce the anti-inflammatory cytokine IL-10 was analyzed. Results in Fig 6J show that stimulated RAW 264.7 cells produced IL-10 in a TLR1/2/4 and NF-κB dependent manner. As opposed to TNF-α, IL-6 and NO induction, the MAPK Kinases pathway appeared to contribute to triggering of IL-10 secretion, as the MEK 1 and MEK 2 inhibitors had a significant effect. All control conditions showed the validity of the *in vitro* experimental setup (Fig 6K and Fig 6L).

## Discussion

Our study provides the comprehensive analysis of virulence factors, antibiotic resistance genes, and determinants involved in interactions of the food-borne *P. agglomerans* KM1 isolate with other microorganisms and eukaryotic hosts. *P. agglomerans* KM1 genome was assembled into one circular chromosome and three mega plasmids. Whole-genome phylogenetic analysis based on ANI values showed that KM1 was 98.68% similar to that of *P. agglomerans* UAEU18, a plant-growth promoting bacterium isolated from date palm rhizosphere soil in the United Arab Emirates [51]. The second closest related stain was *P. agglomerans* C410P1 (98.66%), a biocontrol agent isolated from lettuce in China (GenBank accession number CP016889), followed by *P. agglomerans* Tx10 (97.44%) isolated from the sputum of cystic fibrosis patient [52]. Our KM1 stain was isolated from homemade kimchi, a traditional South Korean fermented side dish prepared using Baechu cabbage (*Brassica rapa* subsp. *pekinensis*) as main ingredient. In this context, we would like to propose that the origin of the KM1 stain is most probably the Baechu cabbage itself as *P. agglomerans* was shown to grow as an epiphyte on vegetables and fruits [68, 69]. However, one cannot exclude contamination of kimchi by humans during the preparation process, as *P. agglomerans* was found in sputum of cystic fibrosis patient. Important to mention here is the fact that the phenotypic analysis of the KM1 strain showed the ability to maintain growth at acidic pH of 4.5 and at 4 °C, the conditions characteristic for kimchi fermentation and storage [3]. In-depth genomic analysis complemented the above information, showing that this Gram-negative bacterium contained a battery of genes conferring resistance to cold and heat shock, as well as genes involved in osmoregulation.

Further genomic characterization excluded the possibility that KM1 is a biocontrol agent, as known biopesticide strains of *P. agglomerans* are capable of producing antibiotics such as pantocins, herbicolins, microcins, and phenazines [14]. In the current study, genome annotation showed that KM1 strain has no genes conferring antibiotic production. Interestingly, the genomic data and results obtained from the Kirby-Bauer disk diffusion method pointed out that KM1 genome carried 13 antibiotic resistance genes conferring resistance to clinically important antibiotics among them penicillin G, bacitracin, vancomycin, rifampicin and fosfomycin. However, genome-based analysis was not able to predict the specific AMR genes that confer resistance to the antibiotics that KM1 was resistant to except for *bacA*, which confers resistance to bacitracin. Recently, Su et al. suggested that these limitations of genome-based prediction arise from scarcity of genetic basis of resistance for some antibiotics [70]. These findings suggest that there is a need for better standardization of AMR annotation to facilitate accurate detection of AMR genes. Previously published clinical reports demonstrated that *P. agglomerans* can be responsible for a wide range of infections encompassing pneumonia, bladder and wound/skin infections, septicemia and meningitis [71]. While bacitracin is an antibiotic prescribed against skin infections, fosfomycin is an effective drug against bladder infections [72-74]. Vancomycin is used to treat meningitis, and rifampicin exerts antimicrobial activity against *Mycobacterium* species, as such being the antibiotic of choice prescribed against tuberculosis [75, 76]. Hence, in the context of our study, these antibiotics would be ineffective against *P. agglomerans* KM1 strain. Previous studies showed that isolates of *P. agglomerans* harbored wide spectrum of antibiotic resistant genes, clinical reports demonstrated cases of pneumonia and death in children with comorbidities where the causative agent was identified as a carbapenem-resistant *P. agglomerans* [12]. Additionally, another study showed that fifty percent of *P. agglomerans* isolates extracted from an infant formula milk were resistant to cefotaxime, moxifloxacin, cotrimoxazole and ticarcillin [10]. As such, the food-borne isolates including *P. agglomerans* KM1 may serve as reservoir of antibiotic resistance genes, posing health risks, food safety concerns, and contributing to the spread of antibiotic resistance through horizontal gene transfer. These findings suggest that *P. agglomerans* may increasingly become more common in clinical settings, as reported in case of major nosocomial pathogens such as *Acinetobacter baumannii* and *Pseudomonas aeruginosa* [77].

The data on the KM1 antibiotic resistance profile prompted us to further investigate the presence of potential virulence factors within the sequenced genome. While in previous studies, most *P. agglomerans* research focused on the characterization of genomic features involved in plant-growth promotion, production of antibiotics, and phenotypic antibiotic susceptibility, only few reports described its genomic determinants of virulence and potential role in pathogenicity. For example, the first genomic characterization of *P. agglomerans* antimicrobial resistance genes isolated from diseased rainbow trout revealed the presence of ARGs *qnrS* and *sul2* [78]. In this context, our study provides new information related to potential virulence factors and pathogenicity of the KM1 isolate. The latter contained putative virulence genes previously identified in *Klebsiella pneumoniae, Escherichia coli*, and *Salmonella enterica*, which are closely related genera of *Pantoea* within the *Enterobacteriaceae* family [14]. Several identified virulence genes deserve special attention in relation to potential pathogenicity such as the genes associated to the type VI secretion system (T6SS), adhesion, iron uptake and sequestration system as well as secretion of toxins such as hemolysins.

Interestingly, the KM1 strain harbors the complete structural gene components of the T6SS as identified using the VFDB and PGAP annotation. It has been reported that the *Hcp* and *vgrG* proteins were considered as hallmarks of a functional T6SS, which target essential cell structures including cell wall, the cell membrane of the genetic material [79, 80]. As such the T6SS was identified in genome sequences of several Gram-negative bacteria and is widely distributed in *Proteobacteria* [81]. In *Pantoea ananatis*, the T6SS complex plays important roles related to its pathogenicity, host range determination, niche adaptation and competition through killing of neighboring bacteria [82]. The T6SS is composed of a membrane complex (*TssJ, TssL* and *TssM)*, a baseplate (*TssK, TssF, TssG, TssE*, and *VgrG*) and a tail complex (*TssB, TssC* and *Hcp*). The hemolysin-coregulated protein *Hcp* and valine-glycine repeat G (*VgrG*) proteins, which are considered the essential components of the T6SS apparatus, were identified in KM1. The *Hcp* is regarded as one of the secreted effectors of T6SS and it forms a tube in the outer component of the T6SS apparatus [83]. Moreover, we also detected the gene encoding the *VgrG* spike protein and PAAR domain-containing protein, which facilitates the translocation of effector proteins to mediate microbe-host interactions [62, 84]. In addition, the ATPase *ClpV* disassembles the contracted tail sheath, which enables a new T6SS complex to be reassembled from the released subunits [63]. Human pathogenic *Pseudomonas aeruginosa* and *Vibrio cholera* were shown to possess T6SS, which functions as a delivery apparatus of bacterial toxins into the host cells [83, 85]. As such, the T6SS is a versatile protein secretion machinery that is able to target eukaryotic cells, highlighting its importance in the context of infection and disease [63]. In *Francisella tularensisis* subsp. *tularensis*, the causative agent of the life-threatening zoonotic disease tularemia, T6SS is essential for entry and multiplication within host macrophages [86]. In case of the isolated KM1 strain, the presence of T6SS may increase its fitness in relation to the host-associated microbial communities or cause pathology to neighboring host cells during infection.

The KM1 strain also harbors genes involved in hemolysis. These genes code for extracellular cytotoxic proteins, which are known virulence factors that target cell membranes causing erythrocyte lysis [87]. The KM1 strain has a *hlyIII* encoding hemolysin III, hemolysin *XhlA*, and hemolysin secretion/activation genes (*Hha, ShlB, FhaC, HecB*). In the adhesion category, the presence of genes encoding filamentous hemagglutinin (*FHA*) and outer membraned protein A (*OmpA*) were detected in KM1 strain. FHA is an important virulence factor that is required for adhesion to the epithelial cells of mammalian hosts [64]. *Bordetella pertussi*s uses FHA as a major adhesion protein to attach itself to the host cells and at the same time increasing the adherence of other pathogens [88]. *OmpA* has essential roles in bacterial adhesion, invasion, and intracellular survival along with evasion of host defenses or stimulators of pro-inflammatory cytokine production [65]. Thus, these findings indicate that *FHA* and *OmpA* in KM1 may have potential pathogenic roles in life-threatening lower respiratory tract infections and urinary tract infections associated to *P. agglomerans* [12]. The last category of identified virulence factors includes genes associated with an iron uptake and siderophore mediated iron sequestration. Iron is an important element for survival and colonization by bacteria since it plays a crucial role in the electron transport chain to produce energy [89]. Iron acquisition systems are used by bacteria to scavenge iron from the environment under iron-restricted conditions [90]. Therefore, successful competition for iron is crucial for pathogenicity. The ferric iron uptake regulator (*Fur*), iron manganese ABC transporter permease (*SitC*) and iron uptake protein (*EfeO*) were all identified in KM1. Ferric uptake regulator (*Fur*) is a transcription factor that upregulates virulence factors in bacteria during iron depletion [91]. These findings suggest the ability of KM1 to survive in the blood and its potential ability to invade the central nervous system by crossing the blood-brain barrier as have been observed in *Cronobacter sakazakii* [92]. Interestingly, the presence of ferric siderophore enterobactin transporter in KM1 may facilitate extraction of iron from host-iron complexes like lactoferrin, transferrin, and hemoglobin [90]. Taken together, these results suggest that siderophore system could be an essential genetic determinant for growth, virulence and potential pathogenicity of *P. agglomerans* KM1.

In order to complete the characterization of isolated *P. agglomerans* KM1, immunostimulatory properties were assessed, using RAW 264.7 macrophages. The heat inactivated whole-cell preparation of *P. agglomerans* triggered secretion of pro-inflammatory cytokines TNF-α and IL-6, as well as NO. At the same time, culture supernatants contained high levels of anti-inflammatory IL-10. While the bacterial cell wall contains lipopolisacharide (LPS), the main immuno-stimulatory molecule and a well-known agonist of TLR4, bacterial triacylated lipoproteins were shown to activate TLR1/2 signaling pathway [93, 94]. Previously published reports attributed TNF-α production mainly to the action of *P. agglomerans* derived LPS, able to trigger activation of the TLR4. Our current study provides an additional evidence that beside the strong involvement of the TLR4, TNF-α section can also be mediated by TLR1/2, however to a lesser extent. It was shown before that binding of TLR4 agonists activates My88/MAL adaptor proteins or triggers endosomal internalization of the receptor/ligand complex. Further internalization leads to interactions with Translocating chain associated membrane protein (TRAM) and TIR domain containing adapter–inducing interferon-β (TRIF) protein resulting in downstream activation of various transcriptional factors. As such, both endosomal and surface activation of the TLR4 signaling can give rise to TNF-α secretion, resulting in NF-kB or MAPKKs activation [95-97]. In this context, our data demonstrated that stimulated RAW 264.7 macrophages secreted TNF-α based on activation of the NF-κB with no involvement of MAPKKs. The dependence on the NF-κB was also demonstrated in case of secretion of IL-6 and NO. Here in contrast to TNF-α, a strong involvement of both TLR1/2 and TLR4 was recorded. Interestingly, the heat inactivated whole-cell preparation of *P. agglomerans* was also capable of inducting IL-10 production. The anti-inflammatory cytokine IL-10 was shown to antagonize the action of pro-inflammatory cytokines such as TNF-α, preventing tissue and cell injury due to induction of an excessive inflammation with cytotoxic activity [98]. The recorded levels of IL-10 were completely reduced to background levels using a RAW 264.7 TLR4 knock-out cell line, showing that IL-10 induction was dependent on TLR4 signaling. The CU-CPT22 inhibitor of the TLR1/2 signaling also significantly reduced IL-10 secretion. However, the effect was only partial as compared to the complete abrogation of IL-10 secretion observed when using the TLR4 knock-out cell line. While the role of TLR2 in section of IL-10 is well documented, the involvement of the TLR4 is less characterized, and as such our study sheds a new light on TLR4 mediated IL-10 secretion [95, 96, 99, 100]. Important to keep in mind is that the abrogation of IL-10 secretion may reflect a cumulative effect of the deletion of the TLR4 present on the surface and TLR4 located in endosomal compartment. As opposed to production of TNF-α, Il-6 and NO production, the secretion of IL-10 involved partial activation of MAPKKs and the dominant role of the NF-κB. The involvement of TLR4 in signaling for IL-10 secretion may have an additional regulatory function in maintaining the balance between secretion of the pro-inflammatory and inflammatory cytokines, controlling at the same time the extent of an injury during inflammation.

In conclusion, the genome analysis of the foodborne isolate *P. agglomerans* KM1 improves our understanding of its virulence determinants indicating potential for pathogenicity. In addition to thirteen antibiotic resistance genes, several virulence factors were identified such as the complete T6SS, filamentous hemagglutinin, siderophore-mediated iron acquisition system (Enterobactin transporter) and hemolysin. The KM1 strain showed strong immuno-stimulatory properties on RAW 264.7 macrophages, with a dominant role of TLR4 signaling and NF-κB activation, resulting in secretion of TNF-α, Il-6, NO and Il-10. Further large-scale studies of clinical isolates are needed to validate the identified virulence factors and antibiotic resistance genes of this potentially opportunistic *P. agglomerans*. Hence, the high quality draft genome of *P. agglomerans* KM1 will provide a baseline for further studies leading to in-depth understanding of molecular mechanisms of *P. agglomerans* pathogenesis.

## Acknowledgment

Preliminary experiments preceding the results presented in this paper were technically executed in part by Miss K. Arora, Miss M. Kim and Mr. N. Vereecke, former members of the laboratory assistant team at Ghent University Global Campus.

## Supporting information

**S1 Fig. Assessment of the completeness of genome assembly and annotation of *P. agglomerans* KM1 using BUSCO**. Colors refer to the percentage of the complete single-copy orthologs (light blue), complete duplicated orthologs (blue), fragmented or incomplete orthologs (yellow), and missing orthologs (red).

**S2 Fig. Phylogenetic relationship of *P. agglomerans* KM1 and its closely related species based on 16S rRNA**. Branch labels represent bootstrap values, in percent, based on 1000 replicates. *Escherichia coli* strain K12 substr. MG1655 was used as outgroup. Accession numbers were reported in the methods section.

**S3 Fig. Functional classifications of *P. agglomerans* KM1 coding sequences against the (A) COG and (B) SEED Subsystem database**.

**S4 Fig. Roary matrix-based protein sequence comparison and the associated phylogenetic tree of *P. agglomerans* strains**.

**S5 Fig. Core and pan-genome analysis of five *P. agglomerans* strains**. (A) Heap’s law chart representation regarding conserved genes and total genes in *P. agglomerans* genomes. (B) Diagram of new genes vs. unique genes in relation to number of genomes embedded in the analysis.

**S6 Fig. Linear genomic map of *P. agglomerans* KM1 phage-associated regions obtained with PHASTER**. Different functional groups of coding sequences of phage origins are denoted by different colors, according to their function.

**S7 Fig. Gel electrophoresis of amplified PCR products of type VI secretion system in *P. agglomerans* KM1**. Lane 1: 100 bp DNA marker, Lane 2: *Hcp* negative control, Lane 3: *Hcp* in KM1, Lane 4: *VgrG* negative control, Lane 5: *VgrG* in KM1. The amplicon size of T6SS effectors were 1301 bp (*Hcp)* and 1011 bp (*VgrG*).

**S1 Table. CRISPR loci of *P. agglomerans* KM1**.

**S2 Table. Genomic islands present in the *P. agglomerans* KM1**.

**S3 Table. Biochemical identification of *P. agglomerans* KM1 using API 20E test kit. S4 Table. Genotypic antibiotic resistance gene profile of *P. agglomerans* KM1**.

## References

1. Park KY, Jeong JK, Lee YE, Daily JW, 3rd. Health benefits of kimchi (Korean fermented vegetables) as a probiotic food. Journal of medicinal food. 2014;17(1):6–20.

2. Patra JK, Das G, Paramithiotis S, Shin HS. Kimchi and other widely consumed traditional fermented foods of Korea: A review. Frontiers in microbiology. 2016;7:1493.

3. Jung JY, Lee SH, Kim JM, Park MS, Bae JW, Hahn Y, et al. Metagenomic analysis of kimchi, a traditional Korean fermented food. Applied and environmental microbiology. 2011;77(7):2264–74.

4. Song WJ, Chung HY, Kang DH, Ha JW. Microbial quality of reduced-sodium napa cabbage kimchi and its processing. Food science & nutrition. 2019;7(2):628–35.

5. Shin J, Yoon KB, Jeon DY, Oh SS, Oh KH, Chung GT, et al. Consecutive outbreaks of Enterotoxigenic *Escherichia coli* O6 in schools in South Korea caused by contamination of fermented vegetable kimchi. Foodborne pathogens and disease. 2016;13(10):535–43.

6. Choi Y, Lee S, Kim HJ, Lee H, Kim S, Lee J, et al. Pathogenic *Escherichia coli* and *Salmonella* can survive in kimchi during fermentation. Journal of food protection. 2018;81(6):942–6.

7. Luziatelli F, Ficca AG, Melini F, Ruzzi M. Genome sequence of the plant growth-promoting rhizobacterium *Pantoea agglomerans* C1. Microbiology resource announcements. 2019;8(44).

8. Humphrey J, Seitz T, Haan T, Ducluzeau AL, Drown DM. Complete genome sequence of *Pantoea agglomerans* TH81, isolated from a permafrost thaw gradient. Microbiology resource announcements. 2019;8(1).

9. Palmer M, de Maayer P, Poulsen M, Steenkamp ET, van Zyl E, Coutinho TA, et al. Draft genome sequences of *Pantoea agglomerans* and *Pantoea vagans* isolates associated with termites. Standards in genomic sciences. 2016;11:23.

10. Mardaneh J, Dallal MM. Isolation, identification and antimicrobial susceptibility of *Pantoea (Enterobacter) agglomerans* isolated from consumed powdered infant formula milk (PIF) in NICU ward: First report from Iran. Iranian journal of microbiology. 2013;5(3):263–7.

11. De Champs C, Le Seaux S, Dubost JJ, Boisgard S, Sauvezie B, Sirot J. Isolation of *Pantoea agglomerans* in two cases of septic monoarthritis after plant thorn and wood sliver injuries. Journal of clinical microbiology. 2000;38(1):460–1.

12. Buyukcam A, Tuncer O, Gur D, Sancak B, Ceyhan M, Cengiz AB, et al. Clinical and microbiological characteristics of *Pantoea agglomerans* infection in children. Journal of infection and public health. 2018;11(3):304–9.

13. Cruz AT, Cazacu AC, Allen CH. *Pantoea agglomerans*, a plant pathogen causing human disease. Journal of clinical microbiology. 2007;45(6):1989–92.

14. Walterson AM, Stavrinides J. *Pantoea*: insights into a highly versatile and diverse genus within the Enterobacteriaceae. FEMS microbiology reviews. 2015;39(6):968–84.

15. Lim JA, Lee DH, Kim BY, Heu S. Draft genome sequence of *Pantoea agglomerans* R190, a producer of antibiotics against phytopathogens and foodborne pathogens. Journal of biotechnology. 2014;188:7–8.

16. Pusey PL, Stockwell VO, Reardon CL, Smits TH, Duffy B. Antibiosis activity of *Pantoea agglomerans* biocontrol strain E325 against *Erwinia amylovora* on apple flower stigmas. Phytopathology. 2011;101(10):1234–41.

17. Shariati JV, Malboobi MA, Tabrizi Z, Tavakol E, Owilia P, Safari M. Comprehensive genomic analysis of a plant growth-promoting rhizobacterium *Pantoea agglomerans* strain P5. Scientific reports. 2017;7(1):15610.

18. Smits TH, Rezzonico F, Kamber T, Blom J, Goesmann A, Ishimaru CA, et al. Metabolic versatility and antibacterial metabolite biosynthesis are distinguishing genomic features of the fire blight antagonist *Pantoea vagans* C9-1. PloS one. 2011;6(7):e22247.

19. Fukasaka M, Asari D, Kiyotoh E, Okazaki A, Gomi Y, Tanimoto T, et al. A lipopolysaccharide from *Pantoea agglomerans* is a promising adjuvant for sublingual vaccinesto induce systemic and mucosal immune responses in mice via TLR4 pathway. PloS one. 2015;10(5):e0126849.

20. Hebishima T, Matsumoto Y, Watanabe G, Soma G, Kohchi C, Taya K, et al. Oral administration of immunopotentiator from *Pantoea agglomerans* 1 (IP-PA1) improves the survival of B16 melanoma-inoculated model mice. Experimental animals. 2011;60(2):101–9.

21. Kobayashi Y, Inagawa H, Kohchi C, Kazumura K, Tsuchiya H, Miwa T, et al. Oral administration of *Pantoea agglomerans*-derived lipopolysaccharide prevents development of atherosclerosis in high-fat diet-fed apoE-deficient mice via ameliorating hyperlipidemia, pro-inflammatory mediators and oxidative responses. PloS one. 2018;13(3):e0195008.

22. Dutkiewicz J, Mackiewicz B, Lemieszek MK, Golec M, Milanowski J. *Pantoea agglomerans*: a marvelous bacterium of evil and good.Part I. Deleterious effects: Dust-borne endotoxins and allergens - focus on cotton dust. Annals of agricultural and environmental medicine : AAEM. 2015;22(4):576–88.

23. Dutkiewicz J, Mackiewicz B, Kinga Lemieszek M, Golec M, Milanowski J. *Pantoea agglomerans*: a mysterious bacterium of evil and good. Part III. Deleterious effects: infections of humans, animals and plants. Annals of agricultural and environmental medicine : AAEM. 2016;23(2):197–205.

24. Tiwari S, Beriha SS. *Pantoea* species causing early onset neonatal sepsis: a case report. Journal of medical case reports. 2015;9:188.

25. Yablon BR, Dantes R, Tsai V, Lim R, Moulton-Meissner H, Arduino M, et al. Outbreak of *Pantoea agglomerans* Bloodstream Infections at an Oncology Clinic-Illinois, 2012-2013. Infection control and hospital epidemiology. 2017;38(3):314–9.

26. Dutkiewicz J, Mackiewicz B, Lemieszek MK, Golec M, Skorska C, Gora-Florek A, et al. *Pantoea agglomerans*: a mysterious bacterium of evil and good. Part II--Deleterious effects:Dust-borne endotoxins and allergens--focus on grain dust, other agricultural dusts and wood dust. Annals of agricultural and environmental medicine : AAEM. 2016;23(1):6–29.

27. Mackiewicz B, Dutkiewicz J, Siwiec J, Kucharczyk T, Siek E, Wojcik-Fatla A, et al. Acute hypersensitivity pneumonitis in woodworkers caused by inhalation of birch dust contaminated with *Pantoea agglomerans* and *Microbacterium barkeri*. Annals of agricultural and environmental medicine : AAEM. 2019;26(4):644–55.

28. Milanowski J, Dutkiewicz J. Increased superoxide anion generation by alveolar macrophages stimulated with antigens associated with organic dusts. Allergologia et immunopathologia. 1990;18(4):211–5.

29. Mudryk M. Plant-isolated *Pantoea agglomerans*--new look into potential pathogenicity. Mikrobiolohichnyi zhurnal. 2012;74(6):53–7.

30. Moghadam F, Couger MB, Russ B, Ramsey R, Hanafy RA, Budd C, et al. Draft genome sequence and detailed analysis of *Pantoea eucrina* strain *Russ* and implication for opportunistic pathogenesis. Genomics data. 2016;10:63–8.

31. Bankevich A, Nurk S, Antipov D, Gurevich AA, Dvorkin M, Kulikov AS, et al. SPAdes: a new genome assembly algorithm and its applications to single-cell sequencing. Journal of computational biology : a journal of computational molecular cell biology. 2012;19(5):455–77.

32. Walker BJ, Abeel T, Shea T, Priest M, Abouelliel A, Sakthikumar S, et al. Pilon: an integrated tool for comprehensive microbial variant detection and genome assembly improvement. PloS one. 2014;9(11):e112963.

33. Gurevich A, Saveliev V, Vyahhi N, Tesler G. QUAST: quality assessment tool for genome assemblies. Bioinformatics. 2013;29(8):1072–5.

34. Simao FA, Waterhouse RM, Ioannidis P, Kriventseva EV, Zdobnov EM. BUSCO: assessing genome assembly and annotation completeness with single-copy orthologs. Bioinformatics. 2015;31(19):3210–2.

35. Tatusova T, DiCuccio M, Badretdin A, Chetvernin V, Nawrocki EP, Zaslavsky L, et al. NCBI prokaryotic genome annotation pipeline. Nucleic acids research. 2016;44(14):6614–24.

36. Lowe TM, Eddy SR. tRNAscan-SE: a program for improved detection of transfer RNA genes in genomic sequence. Nucleic acids research. 1997;25(5):955–64.

37. Lagesen K, Hallin P, Rodland EA, Staerfeldt HH, Rognes T, Ussery DW. RNAmmer: consistent and rapid annotation of ribosomal RNA genes. Nucleic acids research. 2007;35(9):3100–8.

38. Wu S, Zhu Z, Fu L, Niu B, Li W. WebMGA: a customizable web server for fast metagenomic sequence analysis. BMC genomics. 2011;12:444.

39. Aziz RK, Bartels D, Best AA, DeJongh M, Disz T, Edwards RA, et al. The RAST Server: rapid annotations using subsystems technology. BMC genomics. 2008;9:75.

40. Petkau A, Stuart-Edwards M, Stothard P, Van Domselaar G. Interactive microbial genome visualization with GView. Bioinformatics. 2010;26(24):3125–6.

41. Yoon SH, Ha SM, Kwon S, Lim J, Kim Y, Seo H, et al. Introducing EzBioCloud: a taxonomically united database of 16S rRNA gene sequences and whole-genome assemblies. International journal of systematic and evolutionary microbiology. 2017;67(5):1613–7.

42. Kumar S, Stecher G, Li M, Knyaz C, Tamura K. MEGA X: Molecular Evolutionary Genetics Analysis across computing platforms. Molecular biology and evolution. 2018;35(6):1547–9.

43. Darling AC, Mau B, Blattner FR, Perna NT. Mauve: multiple alignment of conserved genomic sequence with rearrangements. Genome research. 2004;14(7):1394–403.

44. Seemann T. Prokka: rapid prokaryotic genome annotation. Bioinformatics. 2014;30(14):2068–9.

45. Page AJ, Cummins CA, Hunt M, Wong VK, Reuter S, Holden MT, et al. Roary: rapid large-scale prokaryote pan genome analysis. Bioinformatics. 2015;31(22):3691–3.

46. Bertelli C, Laird MR, Williams KP, Simon Fraser University Research Computing G, Lau BY, Hoad G, et al. IslandViewer 4: expanded prediction of genomic islands for larger-scale datasets. Nucleic acids research. 2017;45(W1):W30–W5.

47. Arndt D, Grant JR, Marcu A, Sajed T, Pon A, Liang Y, et al. PHASTER: a better, faster version of the PHAST phage search tool. Nucleic acids research. 2016;44(W1):W16–21.

48. Grissa I, Vergnaud G, Pourcel C. CRISPRFinder: a web tool to identify clustered regularly interspaced short palindromic repeats. Nucleic acids research. 2007;35(Web Server issue):W52–7.

49. McArthur AG, Waglechner N, Nizam F, Yan A, Azad MA, Baylay AJ, et al. The comprehensive antibiotic resistance database. Antimicrobial agents and chemotherapy. 2013;57(7):3348–57.

50. Chen L, Yang J, Yu J, Yao Z, Sun L, Shen Y, et al. VFDB: a reference database for bacterial virulence factors. Nucleic acids research. 2005;33(Database issue):D325–8.

51. Alkaabi AS, Sudalaimuthuasari N, Kundu B, AlMaskari RS, Salha Y, Hazzouri KM, et al. Complete genome sequence of the plant growth-promoting bacterium *Pantoea agglomerans* strain UAEU18, isolated from date palm rhizosphere soil in the United Arab Emirates. Microbiology resource announcements. 2020;9(17).

52. Smith DD, Kirzinger MW, Stavrinides J. Draft genome sequence of the antibiotic-producing cystic fibrosis isolate *Pantoea agglomerans* Tx10. Genome announcements. 2013;1(5).

53. Zahradnik J, Plackova M, Palyzova A, Maresova H, Kyslikova E, Kyslik P. Draft genome sequence of *Pantoea agglomerans* JM1, a strain isolated from soil polluted by industrial production of beta-lactam antibiotics that exhibits valacyclovir-like hydrolase activity. Genome announcements. 2017;5(38).

54. Matsuzawa T, Mori K, Kadowaki T, Shimada M, Tashiro K, Kuhara S, et al. Genome sequence of *Pantoea agglomerans* strain IG1. Journal of bacteriology. 2012;194(5):1258–9.

55. Smits TH, Rezzonico F, Kamber T, Goesmann A, Ishimaru CA, Stockwell VO, et al. Genome sequence of the biocontrol agent *Pantoea vagans* strain C9-1. Journal of bacteriology. 2010;192(24):6486–7.

56. Richter M, Rossello-Mora R. Shifting the genomic gold standard for the prokaryotic species definition. Proceedings of the National Academy of Sciences of the United States of America. 2009;106(45):19126–31.

57. Rouli L, Merhej V, Fournier PE, Raoult D. The bacterial pangenome as a new tool for analysing pathogenic bacteria. New microbes and new infections. 2015;7:72–85.

58. Zhang Y, Yan H, Lu W, Li Y, Guo X, Xu B. A novel Omega-class glutathione S-transferase gene in *Apis cerana cerana*: molecular characterisation of GSTO2 and its protective effects in oxidative stress. Cell stress & chaperones. 2013;18(4):503–16.

59. Bible AN, Fletcher SJ, Pelletier DA, Schadt CW, Jawdy SS, Weston DJ, et al. A carotenoid-deficient mutant in *Pantoea* sp. YR343, a bacteria isolated from the rhizosphere of *Populus deltoides*, is defective in root colonization. Frontiers in microbiology. 2016;7:491.

60. Costa E, Usall J, Teixido N, Delgado J, Vinas I. Water activity, temperature, and pH effects on growth of the biocontrol agent *Pantoea agglomerans* CPA-2. Canadian journal of microbiology. 2002;48(12):1082–8.

61. Teixido N, Canamas TP, Abadias M, Usall J, Solsona C, Casals C, et al. Improving low water activity and desiccation tolerance of the biocontrol agent *Pantoea agglomerans* CPA-2 by osmotic treatments. Journal of applied microbiology. 2006;101(4):927–37.

62. Pukatzki S, McAuley SB, Miyata ST. The type VI secretion system: translocation of effectors and effector-domains. Current opinion in microbiology. 2009;12(1):11–7.

63. Ho BT, Dong TG, Mekalanos JJ. A view to a kill: the bacterial type VI secretion system. Cell host & microbe. 2014;15(1):9–21.

64. Inatsuka CS, Julio SM, Cotter PA. *Bordetella* filamentous hemagglutinin plays a critical role in immunomodulation, suggesting a mechanism for host specificity. Proceedings of the National Academy of Sciences of the United States of America. 2005;102(51):18578–83.

65. Confer AW, Ayalew S. The OmpA family of proteins: roles in bacterial pathogenesis and immunity. Veterinary microbiology. 2013;163(3-4):207–22.

66. Haiko J, Westerlund-Wikstrom B. The role of the bacterial flagellum in adhesion and virulence. Biology. 2013;2(4):1242–67.

67. Page MGP. The role of iron and siderophores in infection, and the development of siderophore antibiotics. Clinical infectious diseases : an official publication of the Infectious Diseases Society of America. 2019;69(Suppl 7):S529–S37.

68. Kim D, Hong S, Kim YT, Ryu S, Kim HB, Lee JH. Metagenomic approach to identifying foodborne pathogens on Chinese cabbage. Journal of microbiology and biotechnology. 2018;28(2):227–35.

69. Oie S, Kiyonaga H, Matsuzaka Y, Maeda K, Masuda Y, Tasaka K, et al. Microbial contamination of fruit and vegetables and their disinfection. Biological & pharmaceutical bulletin. 2008;31(10):1902–5.

70. Su M, Satola SW, Read TD. Genome-based prediction of bacterial antibiotic resistance. Journal of clinical microbiology. 2019;57(3).

71. Siwakoti S, Sah R, Rajbhandari RS, Khanal B. *Pantoea agglomerans* infections in children: Report of two cases. Case reports in pediatrics. 2018;2018:4158734.

72. Nguyen R, Khanna NR, Sun Y. Bacitracin topical. StatPearls. Treasure Island (FL) 2020.

73. Bonomo RA, Van Zile PS, Li Q, Shermock KM, McCormick WG, Kohut B. Topical triple-antibiotic ointment as a novel therapeutic choice in wound management and infection prevention: a practical perspective. Expert review of anti-infective therapy. 2007;5(5):773–82.

74. Gardiner BJ, Stewardson AJ, Abbott IJ, Peleg AY. Nitrofurantoin and fosfomycin for resistant urinary tract infections: old drugs for emerging problems. Australian prescriber. 2019;42(1):14–9.

75. Hawley HB, Gump DW. Vancomycin therapy of bacterial meningitis. American journal of diseases of children. 1973;126(2):261–4.

76. Alifano P, Palumbo C, Pasanisi D, Tala A. Rifampicin-resistance, rpoB polymorphism and RNA polymerase genetic engineering. Journal of biotechnology. 2015;202:60–77.

77. Zavascki AP, Carvalhaes CG, Picao RC, Gales AC. Multidrug-resistant *Pseudomonas aeruginosa* and *Acinetobacter baumannii*: resistance mechanisms and implications for therapy. Expert review of anti-infective therapy. 2010;8(1):71–93.

78. Saticioglu IB, Duman M, Altun S. Antimicrobial resistance and molecular characterization of *Pantoea agglomerans* isolated from rainbow trout (*Oncorhynchus mykiss*) fry. Microbial pathogenesis. 2018;119:131–6.

79. Russell AB, Peterson SB, Mougous JD. Type VI secretion system effectors: poisons with a purpose. Nature reviews Microbiology. 2014;12(2):137–48.

80. Durand E, Cambillau C, Cascales E, Journet L. VgrG, Tae, Tle, and beyond: the versatile arsenal of Type VI secretion effectors. Trends in microbiology. 2014;22(9):498–507.

81. Navarro-Garcia F, Ruiz-Perez F, Cataldi A, Larzabal M. Type VI secretion system in pathogenic *Escherichia coli*: Structure, role in virulence, and acquisition. Frontiers in microbiology. 2019;10:1965.

82. Shyntum DY, Venter SN, Moleleki LN, Toth I, Coutinho TA. Comparative genomics of type VI secretion systems in strains of *Pantoea ananatis* from different environments. BMC genomics. 2014;15:163.

83. Mougous JD, Cuff ME, Raunser S, Shen A, Zhou M, Gifford CA, et al. A virulence locus of *Pseudomonas aeruginosa* encodes a protein secretion apparatus. Science. 2006;312(5779):1526–30.

84. Cianfanelli FR, Alcoforado Diniz J, Guo M, De Cesare V, Trost M, Coulthurst SJ. VgrG and PAAR proteins define distinct versions of a functional type VI secretion system. PLoS pathogens. 2016;12(6):e1005735.

85. Pukatzki S, Ma AT, Sturtevant D, Krastins B, Sarracino D, Nelson WC, et al. Identification of a conserved bacterial protein secretion system in *Vibrio cholerae* using the *Dictyostelium* host model system. Proceedings of the National Academy of Sciences of the United States of America. 2006;103(5):1528–33.

86. Clemens DL, Lee BY, Horwitz MA. The *Francisella* type VI secretion system. Frontiers in cellular and infection microbiology. 2018;8:121.

87. Vandenesch F, Lina G, Henry T. *Staphylococcus aureus* hemolysins, bi-component leukocidins, and cytolytic peptides: a redundant arsenal of membrane-damaging virulence factors? Frontiers in cellular and infection microbiology. 2012;2:12.

88. Locht C, Bertin P, Menozzi FD, Renauld G. The filamentous haemagglutinin, a multifaceted adhesion produced by virulent *Bordetella* spp. Molecular microbiology. 1993;9(4):653–60.

89. Krewulak KD, Vogel HJ. Structural biology of bacterial iron uptake. Biochimica et biophysica acta. 2008;1778(9):1781–804.

90. Andrews SC, Robinson AK, Rodriguez-Quinones F. Bacterial iron homeostasis. FEMS microbiology reviews. 2003;27(2-3):215–37.

91. Guo Y, Hu D, Guo J, Li X, Guo J, Wang X, et al. The role of the regulator *Fur* in gene regulation and virulence of *Riemerella anatipestifer* assessed using an unmarked gene deletion system. Frontiers in cellular and infection microbiology. 2017;7:382.

92. Aly MA, Domig KJ, Kneifel W, Reimhult E. Whole genome sequencing-based comparison of food isolates of *Cronobacter sakazakii*. Frontiers in microbiology. 2019;10:1464.

93. Irvine KL, Hopkins LJ, Gangloff M, Bryant CE. The molecular basis for recognition of bacterial ligands at equine TLR2, TLR1 and TLR6. Veterinary research. 2013;44:50.

94. Takeda K, Akira S. Toll-like receptors. Current protocols in immunology. 2015;109:14 2 1–2 0.

95. Li J, Lee DS, Madrenas J. Evolving bacterial envelopes and plasticity of TLR2-dependent responses: Basic research and translational opportunities. Frontiers in immunology. 2013;4:347.

96. Morrison DK. MAP kinase pathways. Cold Spring Harbor perspectives in biology. 2012;4(11).

97. Lawrence T. The nuclear factor NF-kappaB pathway in inflammation. Cold Spring Harbor perspectives in biology. 2009;1(6):a001651.

98. Couper KN, Blount DG, Riley EM. IL-10: the master regulator of immunity to infection. Journal of immunology. 2008;180(9):5771–7.

99. Hu X, Paik PK, Chen J, Yarilina A, Kockeritz L, Lu TT, et al. IFN-gamma suppresses IL-10 production and synergizes with TLR2 by regulating GSK3 and CREB/AP-1 proteins. Immunity. 2006;24(5):563–74.

100. Sanin DE, Prendergast CT, Mountford AP. IL-10 Production in macrophages is regulated by a TLR-driven CREB-mediated mechanism that is linked to genes involved in cell metabolism. Journal of immunology. 2015;195(3):1218–32.

